# Dysregulation of macrophage lipid metabolism underlies intracellular bacterial neuroinvasion

**DOI:** 10.1101/2024.02.28.582462

**Authors:** Zhou Sha, Kun Yang, Shoupeng Fu, Xiaoyong Tong, Hui Yang, Huiling Fang, Qianqian He, Ning Li, Xinyu Shu, Qi Liu, Yongliang Du, Beibei Fu, Yan Xiong, Dong Guo, Jin Liu, Qian Li, Hao Zeng, Xiaokai Zhang, Rui Yao, Xushuo Zhang, Wenjin Guo, Xuhu Mao, Mian Long, Xiaoyuan Lin, Quanming Zou, Haibo Wu

**Affiliations:** School of Life Sciences, Chongqing University, Chongqing 401331, China; Chongqing Public Health Medical Center, Chongqing 400036, China; School of Veterinary Medicine, Jilin University, Changchun 130062, China; School of Pharmaceutical Sciences, Chongqing University, Chongqing 401331, China; Center of Biomechanics and Bioengineering, Key Laboratory of Microgravity (National Microgravity Laboratory), and Beijing Key Laboratory of Engineered Construction and Mechanobiology, Institute of Mechanics, Chinese Academy of Sciences, Beijing 100190, China; School of Engineering Sciences, University of Chinese Academy of Sciences, Beijing 100049, China; Department of Clinical Microbiology and Immunology, College of Pharmacy and Laboratory Medicine, Army Medical University (Third Military Medical University), Chongqing 400038, China; National Engineering Research Center of Immunological, Department of Microbiology and Biochemical Pharmacy, College of Pharmacy and Laboratory Medicine, Army Medical University (Third Military Medical University), Chongqing 400038, China; Department of Pathology, Chongqing Hygeia Hospital, Chongqing 401331, China; Zhejiang Toyouvax Biopharming, Hangzhou 311100, China; Institut für Virologie, Freie Universität Berlin, Robert-von-Ostertag-Str. 7-13, Berlin 14163, Germany

**Keywords:** intracellular bacteria, infection, neuroinvasion, CD36^+^ macrophage, ketogenesis, β-hydroxybutyrate

## Abstract

Acute infection of the central nervous system is one of the most lethal diseases, yet the mechanisms by which intracellular bacteria infiltrate the brain remain poorly understood. Phagocytic cells are typically regarded as the primary site of conflict in the battle against intracellular bacteria; however, little is known about how the intracellular bacteria exploit infected phagocytes to gain access to the brain. In this study, we show that a novel CD36^+^ Fabp4^+^ Pparg^+^ macrophage subpopulation (CD36^+^macrophage) participates in penetration of the brain by intracellular bacteria without disruption of the blood-brain barrier. Biomechanical analysis reveals that the abundance of protrusions and adhesion molecules on CD36^+^ macrophages confers significant resistance to the mechanical stress of blood flow, thereby providing increased opportunities for these macrophages to adhere to the vascular endothelial surface. Metabolomics analysis identifies that macrophage lipid metabolism is dysregulated during bacterial neuroinvasion, and that β-hydroxybutyrate promotes the formation and survival of CD36^+^ macrophages. Importantly, ketogenesis exacerbates symptoms during bacterial neuroinvasion, which could be halted by supplementing with physiological levels of glucose. Collectively, our findings elucidate a pathway by which intracellular bacteria hijack macrophages to invade the brain, suggesting that glycolipid metabolic homeostasis may play a role in the prevention or resolution of bacterial neuroinvasion.

**Highlights:**

1. Intracellular bacteria hijack CD36^+^ Mφ for brain invasion without disrupting the BBB
2. CD36^+^ Mφ has enhanced adhesion to BMECs and increased resistance to shear flow
3. Ketogenesis promotes bacterial neuroinvasion that can be halted by glucose treatment
4. Accumulation of peripheral BHB and CD36^+^ Mφ fuels bacterial neuroinvasion in humans

Graphical Abstract

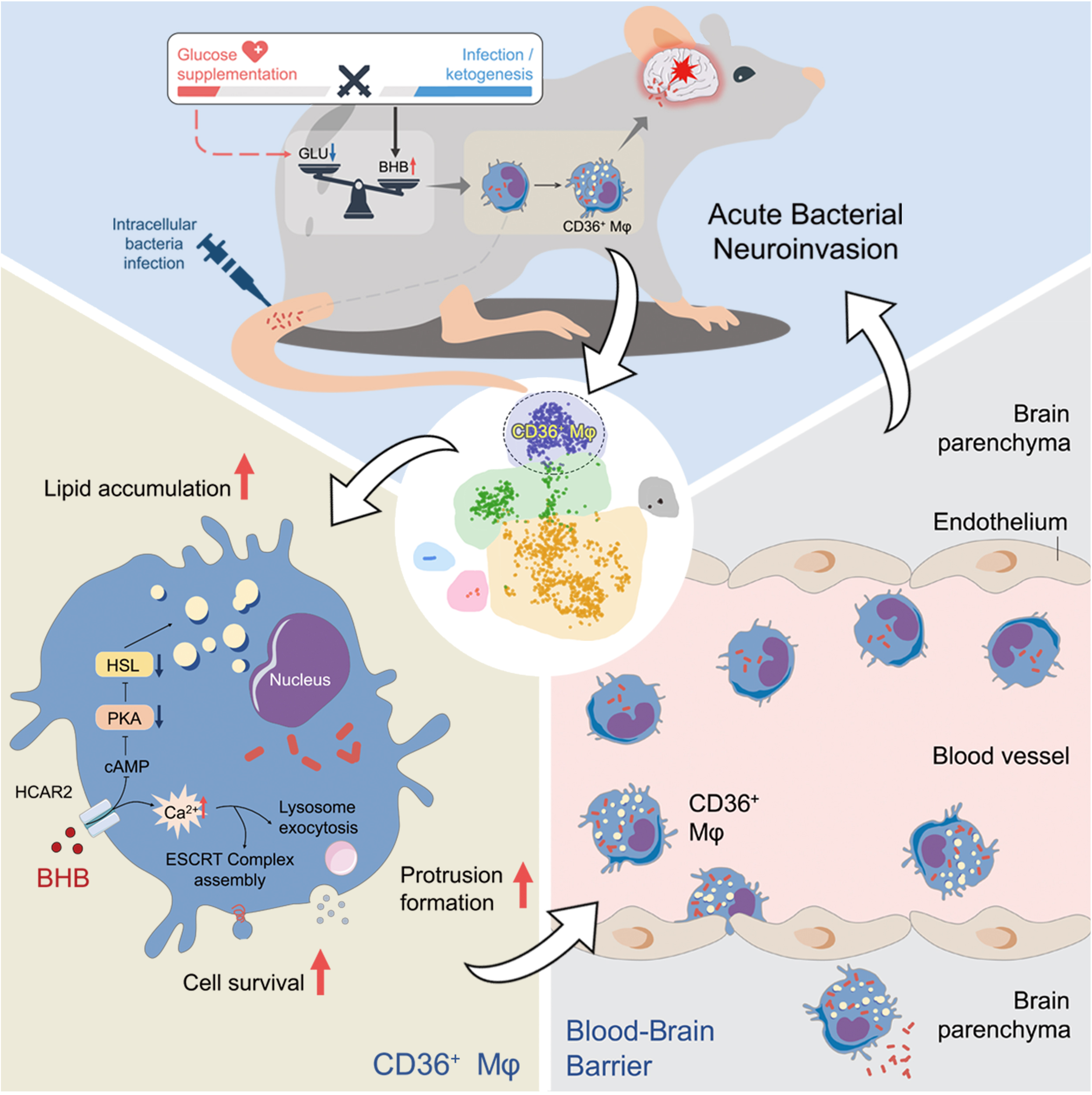

## Introduction

Infections of the central nervous system (CNS) are among the most serious infectious diseases caused by pathogens.^1^ As a life-threatening infection with an extremely high mortality rate, acute bacterial neuroinvasion are usually secondary to infected organs. The blood-brain barrier (BBB) is the foremost line of defense, which consists of brain endothelial cells working in concert with pericytes, smooth muscle cells, astrocytes, microglia and macrophages to protect the CNS from infection.^2^ Currently, pathogens in the bloodstream are thought to penetrate the BBB via three main pathways: transcellular entry, paracellular entry or “Trojan horse” spread.^1^ It is known that some bacteria can enter the brain directly by accumulation during transcytosis or by disruption of tight junctions, accompanied by BBB leakage. For example, *Streptococcus pneumoniae* (*S. pneumoniae*) interacts with host receptors on endothelial cell monolayers to achieve brain invasion by transcytosis.^3–5^ *Neisseria meningitidis* hijacks specific host-pathogen interactions to redistribute tight junctions or adhesion proteins, leading to BBB disruption.^6^ *L. monocytogenes* employs the invasion protein internalin F to facilitate the crossing of the blood-brain barrier ^7,8^, and LLO and ActA have been identified to play a significant role during the invasion of the olfactory epithelium for neurolisteriosis in infants via the intranasal route of infection.^9^ However, the “Trojan horse” route by which the bacteria circumvent the BBB without damaging it has been rarely reported. Efforts to understand the host cells that facilitate the neuroinvasion of intracellular bacteria are urgently needed.

Phagocytes, such as neutrophils and macrophages, play a dominant role in the host response to intracellular bacteria.^10^ The primary function of phagocytes is to engulf and destroy microorganisms and cellular debris through tissue circulation and migration. ^11,12^ By means of phagocytosis and antigen presentation, phagocytes act rapidly as the first line of defense against bacterial infection. However, intracellular bacteria can employ sophisticated strategies to replicate within phagocytes and evade host immune surveillance.^13–15^ Recently, *S. pneumoniae* was reported to hijack receptor activity modifying protein 1 signaling in meningeal macrophages to facilitate brain invasion.^16^ Lecuit et al. showed that the surface protein internalin B of *L. monocytogenes* protects infected monocytes from immune killing by CD8^+^ T cells, thereby prolonging the lifespan of infected cells and facilitating the spread of infection throughout the CNS.^15^ In this context, phagocytes, particularly monocytes and macrophages, may play an important role in neuroinvasion, but how intracellular bacteria take advantage of infected macrophages is not well understood.

The metabolism of host immune cells is reprogrammed during infection. Metabolic signaling, such as glycolysis, helps to awaken the immune cells to resist pathogenic invasion.^17^ In addition, many studies have shown that bacteria manipulate host metabolites to evade clearance by the host immune system.^13,18,19^ *Legionella pneumophila* induces Warburg-like metabolism by secreting the type 4 secretion system effector MitF and decreases oxidative phosphorylation (OXPHOS) in infected macrophages to increase the likelihood of pathogen survival.^18^ A recent report indicates that Nav1.8^+^ nociceptors signal meningeal macrophages via the calcitonin gene-related peptide, and this neuroimmune axis suppresses host defenses and exacerbates bacterial meningitis.^16,19^ Ketogenesis, induced by fasting or dietary supplementation, is a metabolic pathway that provides the body with an alternative form of energy and is closely related to the host immune response.^20,21^ A recent study reported that in pneumonia caused by *Pseudomonas aeruginosa*, ketone bodies are recycled to the lungs via OXPHOS to maintain airway homeostasis.^22^ However, studies have demonstrated that mice fed a ketogenic diet develop a hyperinflammatory response to systemic lipopolysaccharide and an increased mortality rate, which are not attributable to diet-induced microbiota or hyperglycemia.^23^ The role of ketogenesis in the progression and migration of infections remains a topic of contention.^24^ Furthermore, the precise manner in which ketogenesis contributes to the regulation of monocytic/macrophagic homeostasis and the role in bacterial neuroinvasion is poorly defined.

In this study, we discovered a previously unidentified subpopulation of CD36^+^ Fabp4^+^Pparg^+^ macrophages (CD36^+^ macrophages) that serve as immune “Trojan horse” that crosses the BBB without disrupting it, enabling intracellular bacterial neuroinvasion. This subset of macrophages exhibits a foamy-like phenotype, which enhances the ability of the cells to adhere and migrate vertically on the cerebrovascular endothelial surface under the mechanical stress of blood flow. Furthermore, we found that serum BHB, which could be produced by a ketogenic diet, exacerbated bacterial neuroinvasion by increasing the generation and promoting the survival of CD36^+^ macrophages in the peripheral blood. More importantly, we demonstrated that supplementation of mice with physiological levels of glucose rescued the disturbance in lipid metabolism, suppressed the generation of CD36^+^ macrophages, and significantly attenuated the clinical symptoms of encephalitis during bacterial brain invasion. Our study outlines a new common strategy employed by intracellular bacteria to invade the brain, which may be a potential therapeutic target for the treatment of bacterial neuroinvasion.

## Results

### CD36^+^ macrophages are potential carriers of bacteria during brain invasion

To investigate the role of immune cells in bacterial neuroinvasion, we constructed an acute encephalitis mouse model caused by *L. monocytogenes*. This was achieved by testing different infection routes and different doses, including intravenous injection (1×10^5^, 1×10^6^, and 1×10^7^ CFU) (Figure S1A) and intraperitoneal injection (2×10^6^, 2×10^7^, and 2×10^8^ CFU) (Figure S1B). A comprehensive analysis of the survival rate, the bacterial burden in brain tissues, and the pro-inflammatory cytokines in cerebrospinal fluid (CSF) and brain homogenate revealed that both the intravenous injection of 1×10^7^ CFU and the intraperitoneal injection of 2×10^8^ CFU *L. monocytogenes* were successful in establishing an acute neuroinvasion mouse model (Figure S1A and S1B). Additionally, histopathology confirmed the presence of neuronal damage in the cerebral cortices, which is a hallmark of encephalitis (Figure S1C). Behavioral changes, such as diminished activity, paresis, and loss of body weight, were also observed (Figure S1D and S1E). To eliminate the possibility of septic shock resulting from bacteremia, 1×10^7^ CFU of heat-killed and Δhly *L. monocytogenes* were intravenously administered to mice, respectively. The results showed that neither heat-killed nor attenuated *L. monocytogenes* was capable of causing acute neuroinvasion in the same manner as live bacteria, as evidenced by the bacterial loads in the brain, the survival rates, and the clinical scores (Figure S1F-S1H). To further substantiate the hypothesis that intracellular bacteria may utilize phagocytes as a means of gaining access to the brain, gentamicin was administered to mice four hours post-infection with *L. monocytogenes* (Figure S1I). The results showed that when gentamicin was administered prior to infection, there was a substantial remission in blood and brain bacterial loads, as well as an improvement in survival rate of mice (Figure S1I-S1L). However, when gentamicin was administered 4 hours post infection, we found that only the bacteria in the circulation were eradicated, and the brain bacterial loads and survival rate of mice were not significantly different from those of the control group (Figure S1J-S1L). Importantly, Evans blue staining revealed no appreciable extravasation of dye in these *L. monocytogenes*-infected mice (Figure S1M). This finding effectively excludes the possibility that the bacteria gain direct access to the brain by disrupting the BBB. Collectively, these data suggested that intracellular bacteria may exploit host cells to gain access to the brain in acute neuroinvasion.

To identify the specific carrier in the acute bacterial neuroinvasion model, peripheral blood mononuclear cells (PBMCs) were collected and single-cell RNA sequencing (scRNA-seq) was performed (Figure 1A). Immune cell lineages were identified using canonical surface markers of dendritic cells (DCs: *Xcr1, Flt3, CD209a*), NK cells (*Nkg7*), T cells (*CD3*), monocyte/macrophages (Mo/Mφ: *Fcgr1, Adgre1*), B cells (*CD79*), neutrophils (*S100a9, S100a8*; Figure 1B and 1C).^25^ Among these lineages, Mo/Mφ is involved in phagocytosis and intracellular killing of microorganisms; however, intracellular bacteria can also reside within these cells, thereby establishing a long-term latent infection.^26^ In this case, Mo/Mφ (n=3535 cells) were extracted and reclustered *in silico* to precisely delineate subpopulations and their distributions before and after neuroinvasion. Based on differentially expressed cell surface markers and transcripts, we annotated Mo/Mφ into 6 subpopulations (Figure 1D, S2A). The results showed that the percentage of Cluster 2 cells was sharply increased in the neurolisteriosis group (Figure S2B). Further, we found that Cluster 2 cells expressed high levels of genes related to lipid accumulation, including *CD36*, peroxisome proliferator-activated receptor gamma (*Pparg*), fatty-acid binding protein 4 (*Fabp4*), *CD9*, apolipoprotein E epsilon (*ApoE*), and triggering receptor expressed on myeloid cells 2 (*Trem2*; Figure 1E).^27–31^

**Figure 1.**
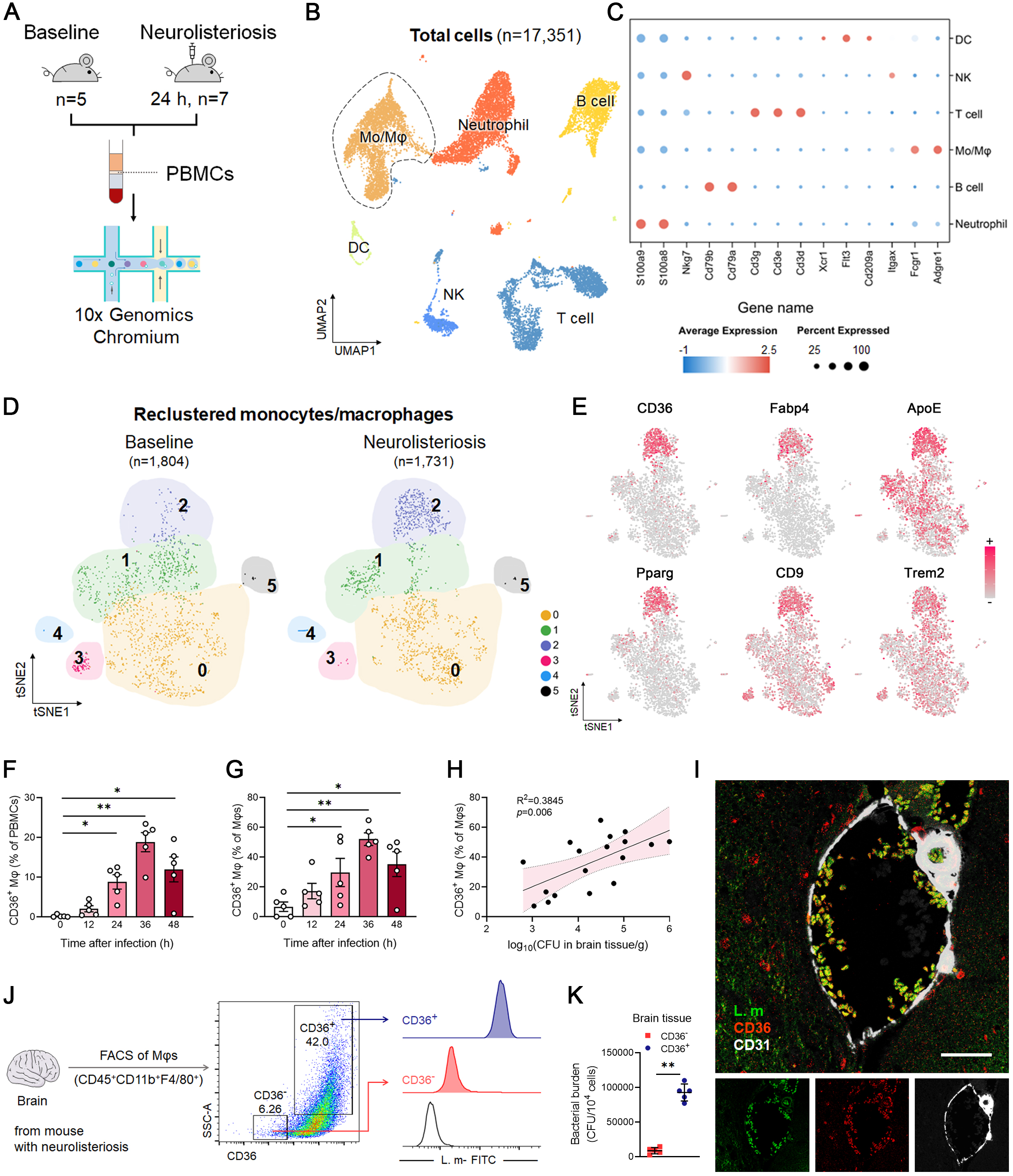
Identification of CD36^+^ macrophage during neurolisteriosis. (A) Summary of the experimental design (control mice, n=5; neurolisteriosis mice, n=7). (B) Uniform Manifold Approximation and Projection (UMAP) representation of integrated transcriptome-based clustering. A total of 17,351 peripheral blood mononuclear cells (PBMCs) were clustered and major immune cell lineages were identified. (C) Dot plot showing the expression of selected transcripts in immune cell clusters. All transcripts were significantly enriched in the pertinent clusters with adjusted p < 0.05. (D) t-distributed Stochastic Neighbor Embedding (tSNE) analysis of the re-clustered cells corresponding to monocytes and macrophages (baseline, n=1,804; neurolisteriosis, n=1,731). (E) Expression of the indicated transcripts (encoding genes associated with lipid accumulation) was projected onto a monocyte/macrophage tSNE plot. (F-G) Mice was intraperitoneally injected with 2×10^7^ CFU *L. monocytogenes* (n=5 for each group). Flow cytometry was performed to assess the percentages of CD36^+^ macrophages in PBMCs (F) and total macrophages (G). (H) Correlation between CD36^+^ macrophages and bacterial burdens in the brain during neurolisteriosis. R, correlation coefficient. (I) Immunofluorescence staining of infected CD36^+^ macrophages in cerebral vasculature of mice with neurolisteriosis. Infected CD36^+^ macrophages were stained with CD36 (red) and *L. monocytogenes* (green), while vascular endothelial cells were stained with CD31 (white). Scale bar, 100 μm. (J-K) FACS of CD36^-^ and CD36^+^ macrophages in the brain tissues (J). The intracellular bacterial burdens were evaluated by flow cytometry (J) and plate dilution method (K). Experiments were performed in duplicate (F-K). Data are presented as mean ± SEM [n=3 in (K), n=5 in (F, G)]. *p < 0.05, **p < 0.01. Abbreviations: PBMCs, peripheral blood mononuclear cells; CD36^+^ Mφs, CD36^+^ macrophages; CFU, colony forming unit.

In order to gain insight into the characteristics of Cluster 2 macrophages, CD36^+^ and CD36^−^ macrophages were isolated from peripheral blood by fluorescence-activated cell sorting. In comparison to CD36^−^ macrophages, the CD36^+^ subpopulation exhibited a markedly elevated level of granularity (SSC^high^; Figure S2C), indicative of an increased number of proteins or organelles within their intracellular compartment and the presence of protrusions on their surfaces. The sorted macrophages were cultured *in vitro* overnight and then labeled with BODIPY FL, a lipophilic probe for polar and neutral lipids. The results showed that the CD36^+^ subpopulation exhibited a significantly greater accumulation of lipid droplets than the CD36^−^ macrophages (Figure S2D). Furthermore, Oil Red O staining was conducted to corroborate this finding (Figure S2D). To confirm the lipid uptake ability of the CD36^+^ macrophages, we employed the use of fluorescently labeled 1,1′-dioctadecyl-3,3,3′,3′-tetramethylindocarbocyanine perchlorate (Dil)-conjugated oxidized low-density lipoprotein (ox-LDL), and the uptake of Dil-conjugated ox-LDL particles by CD36^+^ and CD36^-^ macrophages was quantified. Our findings revealed that CD36^+^ macrophages internalized more than twice the amount of ox-LDL compared to CD36^−^macrophages (Figure S2E). In addition to CD36, other scavenger receptors, including MARCO, SRA1, LDLR and MSR1, have also been implicated in the accumulation of lipids.^32^ Consequently, qRT-PCR assays were performed in order to examine the expression of these receptors in CD36^+^ macrophages. The results demonstrated that only the expression of LDLR was significantly increased in this subpopulation (Figure S2F), a finding that is consistent with the scRNA-seq data.

Subsequently, flow cytometry was performed to ascertain the percentage alterations in CD36^+^ macrophages over time in the context of neurolisteriosis (Figure S2C). The results showed that the percentage and absolute number of CD36^+^ macrophages in the peripheral blood increased gradually over time during neuroinvasion (Figure 1F, S2G), with the CD36^+^ subgroup constituting more than 52% of all macrophages (Figure 1G). Further, the CD36^+^ subpopulation demonstrated the capacity to harbor a greater quantity of bacteria (Figure S2H and S2I), and the percentage of CD36^+^ macrophages in peripheral blood was observed to be positively correlated with the bacterial load in the brain (Figure 1H). These data implied that CD36^+^ macrophages may be involved in bacterial neuroinvasion. In addition, immunofluorescence co-localization analysis was used to validate this hypothesis. The results showed that CD36^+^ macrophages attached to brain microvascular endothelial cells (BMECs) increased with the progression of neuroinvasion in the neurolisteriosis mice (Figure S2J). Further, transection of cerebral blood vessels was performed, and the vascular endothelial cells were identified using an anti-CD31 antibody. The results showed that a substantial number of CD36^+^ macrophages and *L. monocytogenes* co-localized and aggregated on the surface of the vascular endothelial cells (Figure 1I). In addition, CD36^+^ and CD36^−^ macrophages were isolated from brain tissues of the neurolisteriosis mice. The results obtained by flow cytometry, CFU counting, and immunofluorescence indicated that the *L. monocytogenes* was located in the CD36^+^ macrophages rather than CD36^-^ macrophages (Figure 1J and 1K, S2K and S2L). As there was a clear inflammatory response in the brain of mice during neuroinvasion, we subsequently examined the inflammatory profile of the CD36^+^ macrophages. The results showed that CD36^+^ macrophage subpopulation isolated from mice exhibited an anti-inflammatory feature (Figure S2M), suggesting that the inflammatory phenotype of the mice brain is not dependent on the characteristics of CD36^+^ macrophages themselves. Taken together, these results suggested that CD36^+^ macrophages may play a pivotal role in bacterial brain invasion.

### CD36^+^ macrophages facilitate transport of intracellular bacteria across the blood-brain barrier

Ox-LDL has been reported to induce a foamy-like phenotype by enhancing CD36 expression.^33^ Consequently, to obtain CD36^+^ macrophages in large quantities *in vitro* in a rapid and efficient manner, we developed a method to induce the conversion of peritoneal macrophages to CD36^+^ macrophages by stimulation with different concentrations of ox-LDL for 24 hours (Figure S3A). To investigate whether *in vitro* generated CD36^+^ macrophages are analogous to those isolated from peripheral blood, the purified peritoneal macrophages from mice were treated with ox-LDL (80 μg/mL) and then CD36^+^ macrophages were sorted with an APC-conjugated CD36 antibody (Figure S3B). Through Western blotting and RNA-seq analysis, we found that CD36^+^macrophages generated by ox-LDL treatment *in vitro* exhibited Fabp4 and Pparg expression patterns (Figure S3C), and transcriptomic profiling that were highly similar to those of CD36^+^ macrophages isolated from peripheral blood (Figure S3D and S3E). Furthermore, we showed that ox-LDL treatment influenced the development of CD36^+^ macrophages (Figure S3A), however this signal did not impact the inflammatory profile once CD36^+^ macrophages had been formed (Figure S3F and S3G).

Next, to ascertain the role of CD36^+^ macrophages in bacterial neuroinvasion, we constructed an *in vitro* BBB model comprising mouse primary BMECs and neurons in a Transwell system (Figure 2A). The integrity of the cell monolayer was evaluated by measuring the transepithelial electrical resistance (TEER) values using a Millicell-ERS voltohmmeter, and cell monolayers with TEER values above 300 Ω/cm^2^ were considered intact and functional.^34^ We showed that the integrity of the *in vitro* BBB model was not significantly affected by high dose and short period of bacterial infection (Figure S3H). Subsequently, macrophages were infected with *L. monocytogenes* (MOI=10) and then inoculated into the upper Transwell chamber of the BBB model. The transformation of peritoneal macrophages into CD36^+^ macrophages was confirmed to result in an increase in the intracellular bacterial load (Figure S3I). Additionally, the assistance of ox-LDL-stimulated macrophages was observed to significantly increase the number of *L. monocytogenes* organisms crossing the BBB in a dose-dependent manner (Figure 2B). Furthermore, crystal violet staining also demonstrated a dose-dependent increase in the number of macrophages penetrating the endothelial monolayer upon stimulation with ox-LDL (Figure 2C and 2D). To validate whether *L. monocytogenes* reside in the CD36^+^ macrophages and cross the BBB, bacterial staining of these penetrating macrophages was performed. The results showed that bacteria were present in the cells and successfully crossed the BBB in the *in vitro* model (Figure S3J and S3K). In order to conduct further investigation into the capacity of the CD36^-^ but with high lipid content subpopulation (CD36^-^ lipid^high^) to promote neuroinvasion, we tested whether these CD36^-^ lipid^high^ macrophages could facilitate bacterial crossing of the barrier using an *in vitro* BBB model. Interestingly, we did not observe significant differences in the bacterial burden between CD36^-^ lipid^high^ and CD36^-^ lipid^low^ macrophages (Figure S3L-S3N). This result suggested that, despite the crucial role of lipid metabolism in the formation and function of CD36^+^ macrophages, the CD36 receptor may have a hitherto unknown function during the bacterial neuroinvasion.

**Figure 2.**
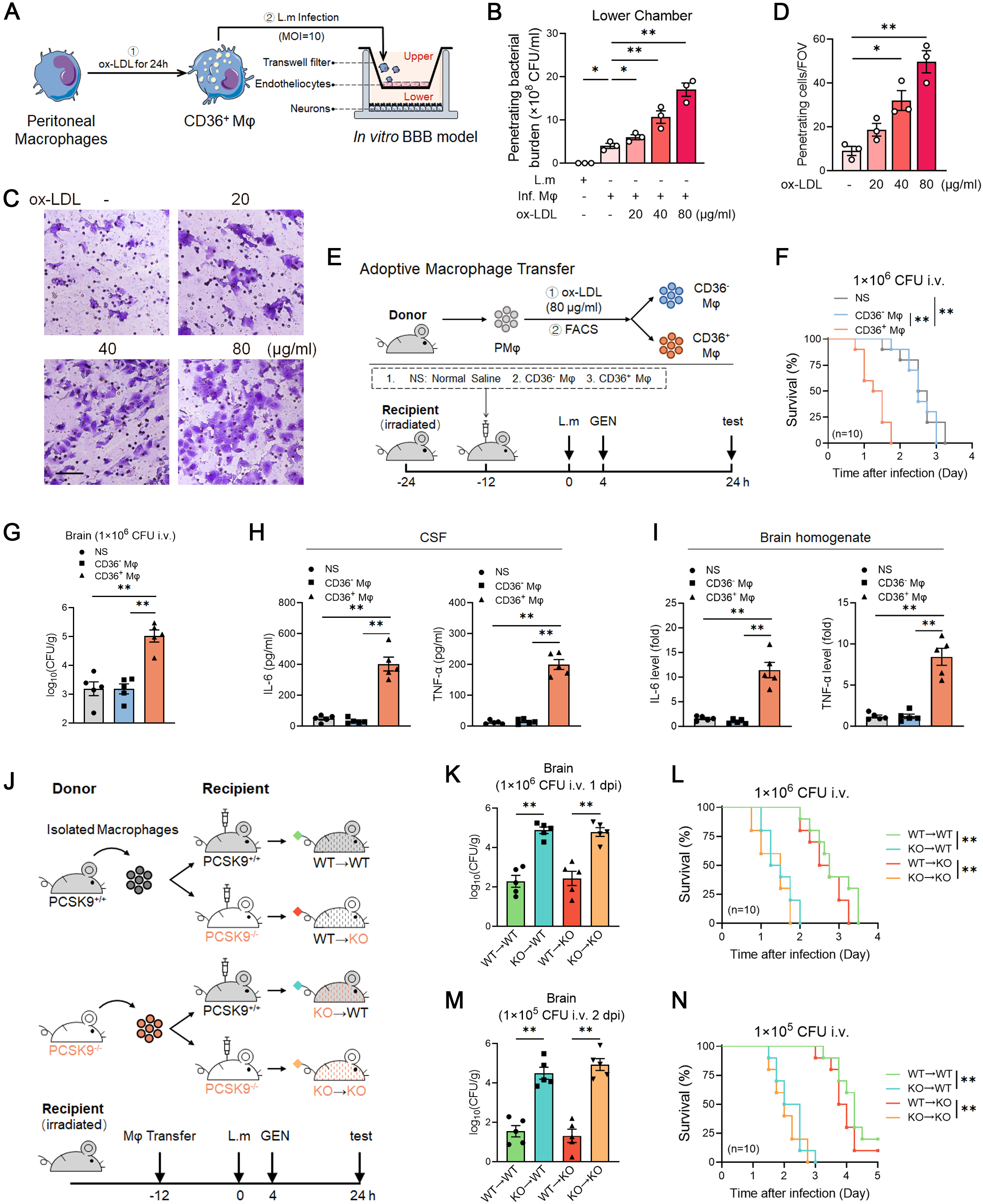
Infected CD36^+^ macrophages transfer bacteria into brain. (A-D) (A) Schematic diagram of the *in vitro* BBB model based on Transwell system. Macrophages were exposed to *L. monocytogenes* (MOI=10) for 6 hours and then treated with 100 µg/ml gentamicin for 2 hours before being added into the upper chamber. After 12 hours of penetration, neuron cells in the lower chamber were lysed for the quantification of penetrating bacteria. (B) Quantification of penetrating bacterial burden in the neuron cells in the lower chamber. (C) Representative images of macrophages penetrating into the lower chamber. Scale bar, 100 μm. (D) Quantification of the penetrating cells in each field of view. (E-I) (E) Experimental diagram of adoptive macrophage transfer. Recipient mice were anesthetized for irradiation (Co 60, 10 Gy). Twelve hours later, adoptive macrophages transfer was performed. After recovery for another 12 hours, recipient mice were intravenously injected with 1×10^6^ CFU of *L. monocytogenes.* Gentamicin (30 mg/kg) was administered intraperitoneally 4 hours after infection. Samples were collected for examination at 24 hours post infection. (n=10 for each group) (F) Survival of recipient mice. (G) Bacterial burdens in brains. (H-I) Pro-inflammatory factor levels in CSF (H) and brain homogenates (I) of recipient mice. (J-N) (J) Schematic diagram of adoptive macrophage transfer experiments in PCSK9^+/+^ (WT) and PCSK9^-/-^ (KO) mice. Recipient mice were irradiated and adoptive macrophages transfer was performed, and then mice were intravenously infected with 1×10^6^ CFU (K-L) or 1×10^5^ CFU (M-N) of *L. monocytogenes*. Gentamicin was administered intraperitoneally 4 hours after infection. (n=10 for each group) (K, M) Bacterial burdens in the brains of recipient mice after intravenous injection of 1×10^6^ CFU *L. monocytogenes* 1 day post infection (K) and 1×10^5^ CFU *L. monocytogenes* 2 days post infection (M). (L, N) Survival of recipient mice after intravenous injection of 1×10^6^ CFU (L) and 1×10^5^ CFU (N) *L. monocytogenes*. Experiments were performed in duplicate (E-N) or triplicate (A-D). Data are presented as mean ± SEM [n=3 in (B, D), n=5 in (G-I, K, M)]. *p < 0.05, **p < 0.01. Abbreviations: ox-LDL, oxidized low density lipoprotein; FOV, field of view; NS, normal saline; PMφ, peritoneal macrophages; CD36^-^ Mφ, CD36^-^ macrophages; CD36^+^ Mφ, CD36^+^ macrophages; i.v., intravenously; i.p., intraperitoneally; ns, nonsignificant.

To ascertain whether the rise in CD36^+^ macrophages facilitated the transfer of *L. monocytogenes* into the brain *in vivo*, we performed adoptive macrophage transfer experiments via tail vein injection based on an irradiated mouse model (Figure 2E). Bone marrow irradiation-based immunosuppression resulted in mice being more susceptible to bacterial infection, which necessitated a reduction of the infectious dose to 1×10^6^ CFU of *L. monocytogenes.* The results showed that the survival time was shortened, the body weight was reduced, and the comprehensive clinical score was increased in the mice receiving CD36^+^ macrophages compared with the control group (Figure 2F, S3O and S3P). Furthermore, we showed that the brain bacterial loads in mice receiving CD36^+^ macrophages were significantly higher than those in mice receiving CD36^-^ macrophages (Figure 2G). Interestingly, no significant differences were observed in bacterial loads among the lung, spleen, and liver (Figure S3Q).

Additionally, we examined the levels of pro-inflammatory factors in the CSF and brain homogenate. The results showed a notable elevation in interleukin 6 (IL-6) and tumor necrosis factor alpha (TNF-α) levels (Figure 2H and 2I), indicating a more severe encephalitis phenotype in the CD36^+^ macrophage transfer group. Importantly, the brains of mice transferred with CD36^+^ macrophages exhibited no discernible Evans blue extravasation, suggesting that these macrophages facilitated the transfer of *L. monocytogenes* into the brain without disrupting the BBB (Figure S3R and S3S).

The results of the pathological histology analysis confirmed the existence of more severe inflammation and neuronal damage in the cerebral cortices of the mice transferred with CD36^+^ macrophages than in the control group (Figure S3R and S3T). Furthermore, when mice were infected with a relatively low dose (1×10^5^ CFU/mouse) of *L. monocytogenes*, the difference in bacterial load in brain tissues between the CD36^+^ macrophage transferred mice and the control group was also observed at day 2 post infection (Figure S3U-S3W). These data suggest that neuroinvasion mediated by CD36^+^ macrophages is not limited to an acute infection model induced by a high dose of bacteria.

We subsequently investigated whether CD36^+^ macrophages may perform a comparable function during the invasion of the brain by other intracellular bacteria. In the pertinent experiments, a positive correlation was observed between CD36^+^ macrophages and neuroinvasion in mice infection models of the intracellular bacterium *Salmonella typhimurium* (*S. typhimurium*; Figure S4A and S4B). The same results were observed in an infection model of Methicillin-resistant *Staphylococcus aureus* (MRSA; Figure S4C and S4D), which, although not a classical intracellular pathogen, exhibits significant intracellular habitation.^35^ The results of the immunofluorescence and bacterial burden analyses of the macrophages isolated from PBMCs of infected mice showed that *S. typhimurium* and MRSA were predominantly present in the CD36^+^ macrophages (Figure S4E and S4F). However, this finding was not validated in a neuroinvasion model of the extracellular bacterium *S. pneumoniae* (Figure S4G and S4H). Next, adoptive macrophage transfer experiments were conducted to further substantiate the hypothesis that CD36^+^ macrophages serve as carriers in the intracellular bacterial invasion of the brain. Once more, disparities in brain tissue bacterial loads were discerned between the CD36^+^ macrophage-transferred and control groups in the neuroinvasion model of MRSA (Fig. S4I-S4K), though not in the model utilizing *S. pneumoniae* (Figure S4L-S4N). Taken together, these data suggested that CD36^+^ macrophages contributed to the process of intracellular bacterial neuroinvasion.

Proprotein convertase subtilisin/kexin 9 (PCSK9) belongs to the proprotein convertase family, members of which participate in cholesterol and fatty acid (FA) metabolism.^36,37^ The intracellular accumulation of low-density lipoprotein cholesterol (LDL-C) in PCSK9^-/-^ is attributed to the inability of the LDL receptor, which is located on the cell membrane, to undergo normal degradation.^38,39^ The mRNA and protein levels of PCSK9 in CD36^+^ macrophage subsets promoted by *L. monocytogenes* infection and in peritoneal macrophages exposed to ox-LDL were examined. The results showed that PCSK9 levels were significantly reduced in both circulating and oxLDL-induced CD36^+^ macrophages (Figure S5A and S5B). Further, we found that macrophages lacking PCSK9 exhibited markedly elevated lipid content in comparison to the control group (Figure S5C), a finding that was corroborated by the expression of FABP4 and PPARG (Figure S5D). Also, it was observed that a greater number of *L. monocytogene* organisms were able to parasitize PCSK9^-/-^ macrophages than those parasitizing wild-type (WT) cells (Figure S5E). In the context of neurolisteriosis, PCSK9^-/-^ mice exhibited a higher percentage of CD36^+^ macrophages in peritoneal blood than their littermates (Figure S5F). In addition, the results of survival rate, bacterial burden in brain tissues, and pro-inflammatory cytokines in CSF revealed that PCSK9^-/-^ mice developed a more pronounced bacterial neuroinvasion in comparison to their littermates following tail vein injection of 1×10^7^ CFU of *L. monocytogenes* (Figure S5G-S5I). Consequently, PCSK9^-/-^ mice were utilized as a model for the spontaneous formation of CD36^+^ macrophages, and the veracity of our findings was validated through adoptive macrophage transfer. Subsequent to the irradiation and a 12-hour recovery period post transfer, recipient mice were intravenously inoculated with 1×10^6^ CFU of *L. monocytogenes*. Gentamicin was used to eliminate extracellular bacteria. Two days later, samples were collected for further examination (Figure 2J). We found that the brain bacterial loads were significantly higher in PCSK9^+/+^ recipients transferred with PCSK9^-/-^ macrophages than those that receiving WT macrophages (Figure 2K). Similar results were obtained in the PCSK9^-/-^ recipients (Figure 2K). However, the differences in bacterial were observed exclusively in the brain tissues, and not in other organs such as the lung, spleen, and liver (Figure S5J). Consistently, the mortality rate, and pro-inflammatory factors in the CSF and brain homogenate were exacerbated in both PCSK9^+/+^ and PCSK9^-/-^ recipients by the transfer of PCSK9^-/-^ macrophages (Figure 2L, S5K and S5L). We further confirmed the aforementioned finding by intravenous injection of 1×10^5^ CFU of *L. monocytogenes* in the same experimental model (Figure 2M and 2N, S5M-S5O).

Furthermore, in order to test the impact of PCSK9 function on the pathology driven by CD36^+^ macrophages during *L. monocytogenes* infection, adoptive macrophage transfer of CD36^+^ and CD36^-^ macrophages sorted from the PCSK9^-/-^ mice were performed (Figure S5P). The results showed that the brain bacterial loads were significantly higher in recipients receiving PCSK9^-/-^CD36^+^ macrophages compared with those receiving PCSK9^-/-^CD36^-^ macrophages (Figure S5Q and S5R). This finding was further corroborated by elevated mortality rates and pro-inflammatory factors in the CSF (Figure S5S and S5T). These results suggested that, while PCSK9 regulates CD36^+^ macrophage formation by modulating intracellular lipids, it does not function directly in CD36^+^ macrophage-driven pathology during bacterial neuroinvasion. Taken together, these data indicated that excessive lipid accumulation contributed to the formation of CD36^+^ macrophages, thereby facilitating neuroinvasion by intracellular bacteria.

### Shear flow analysis demonstrates increased adhesion and resistance of CD36^+^ macrophages during neuroinvasion

We next asked how CD36^+^ macrophages exploited their properties to accumulate on the cerebral blood vessel wall and participate in the process of intracellular bacterial neuroinvasion. Using a two-dimensional live-cell flow chamber assay with a microfluidic chip^40^, we deciphered the multistep process of how macrophages adhered to and moved on the surface of vascular endothelium under physiological flow (Figure 3A). Cellular adhesion ability was calculated by counting total macrophages adhering to the endothelial monolayer under a given shear stress. We found that an increase in the percentage of CD36^+^ macrophages was concomitant with an increase in the number of cells adhering to the endothelial monolayer (Figure 3B and 3C).

**Figure. 3.**
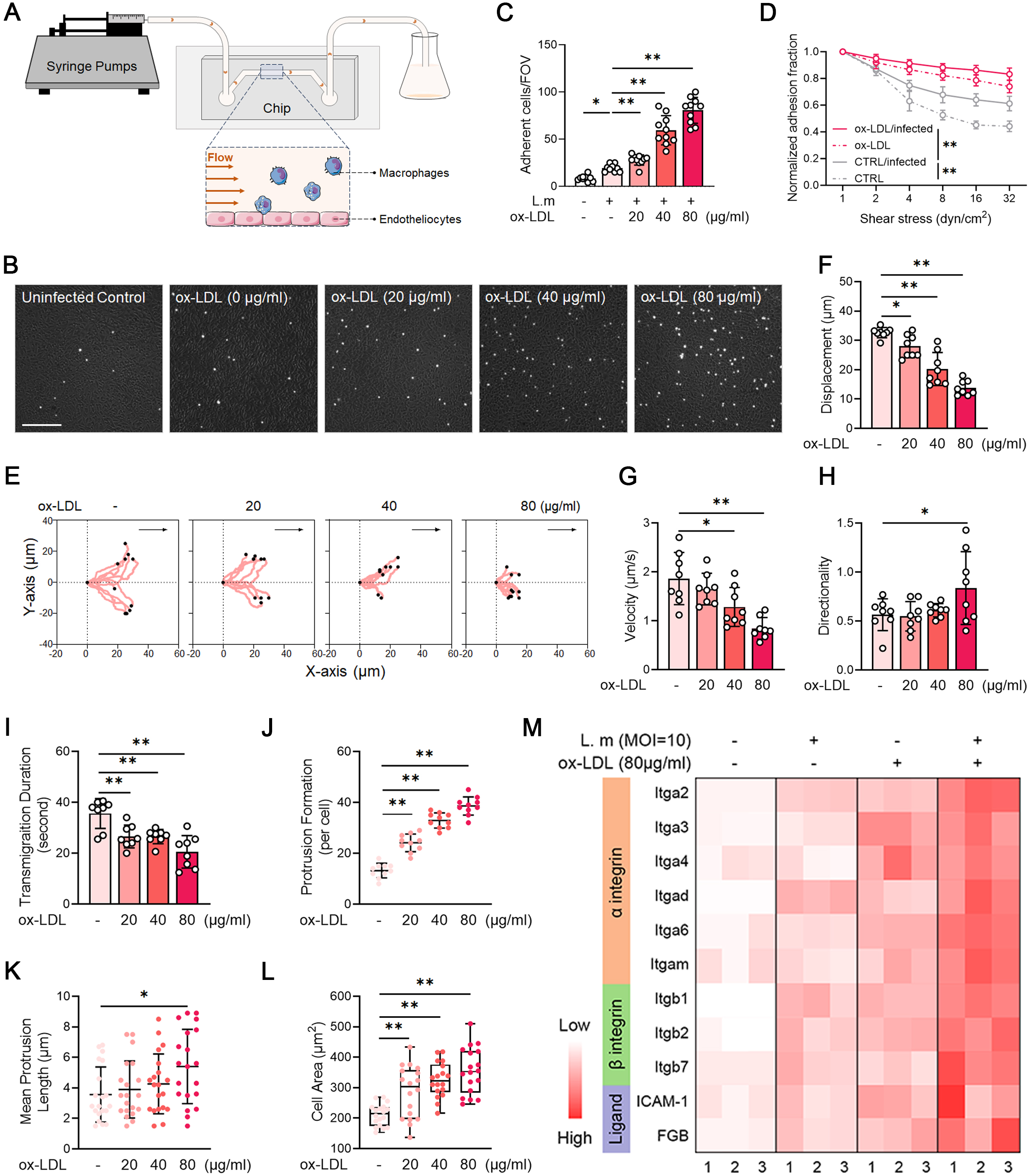
Characterization of infected CD36^+^ macrophages. (A) Schematic diagram of live-cell flow chamber experiment. Primary peritoneal macrophages were pretreated with ox-LDL for 24 hours and then infected with *L. monocytogenes* (MOI=10) for 6 hours. Then cells were perfused over pre-formed endothelial monolayer for 8 minutes at a shear stress of 0.1 dyn/cm^2^ and recorded at 2 dyn/cm^2^ for 10 minutes unless otherwise indicated. (B-C) (B) Optical images of ox-LDL pretreated macrophages adhering to the surface of monolayer endothelial cells under shear flow. Scale bar, 200 μm. (C) Quantification of adherent cells in each field of view. (D) Macrophages resistance to shear flow after adhesion to endothelial cells. Peritoneal macrophages were pretreated with ox-LDL for 24 hours and then challenged with *L. monocytogenes* (MOI=10) for 6 hours. Upon 10-minute adherence of equal macrophages to the monolayer endothelial cells at 1 dyn/cm^2^, the shear stress was stepwise increased to 2, 4, 8, 16, and 32 dyn/cm^2^ for 30 seconds sequentially to detach those adhered cells. (E-H) Crawling dynamics of ox-LDL pretreated macrophages under shear flow. (E) Typical trajectories of ox-LDL pretreated macrophages crawling on monolayer endothelial cells. For each trajectory, the arrested sites of macrophages were overlaid at the origin of the diagram (x/y=0/0) and the corresponding end points were indicated with dots. Arrows indicate the flow direction. (F-H) Quantification of crawling displacement (F), velocity (G) and directionality (H). (I) Transmigration duration of ox-LDL pretreated macrophages after adhesion to the surface of monolayer endothelial cells. (J-K) Quantification of the number (J) and mean length (K) of protrusions in the macrophages pretreated with ox-LDL. (L) Quantification of cell area in the macrophages pretreated with ox-LDL. (M) qRT-PCR analysis of relative expression of genes associated with intercellular adhesion molecules. Experiments were performed in triplicate. Data are presented as mean ± SEM [n=8 in (C, I), n=10 in (F-H), n=18 in (L), n=20 in (K)]. *p < 0.05, **p < 0.01. Abbreviations: ox-LDL, oxidized low density lipoprotein.

Immunofluorescence was performed to show that *L. monocytogenes* were residing in the CD36^+^ macrophages in the biomechanical analysis (Figure S6A). Because adherent cells could be detached by an increase in shear stress, we tested the dynamics of CD36^+^ macrophages’ shear resistance. The results showed that such resistance was significantly stronger than that of control macrophages: CD36^+^ macrophages generated either from ox-LDL treatment or bacterial infection, exhibited remarkable resistance to high shear stress (Figure 3D). Further, we investigated the effect of endothelial density on macrophagic adhesion. The results showed that endothelial cell density, rather than cell type, greatly affected macrophagic adhesion, and CD36^+^ macrophages tended to adhere to the surface of dense endothelial cells (Figure S6B-S6E).

Once attached to the endothelial monolayer, the macrophage crawls along it and finds a place to cross it in the presence of blood flow.^41,42^ We next tested the performance of CD36^+^ macrophages in post-arrest crawling on the endothelial monolayer. We acquired sequential images of macrophages at a shear stress of 2 dyn/cm^2^ and then analyzed them by defining various parameters of crawling dynamics (Figure S6F). The observations were presented by crawling trajectories of each individual cells (Figure 3E), and we found that the crawling ability of adherent CD36^+^ macrophages was dampened, reflecting in shortened crawling displacement and slower crawling speed (Figure 3F and 3G). However, we saw no obvious differences in crawling duration (Figure S6G). Directionality is also a critical aspect of cell crawling dynamics, which is quantified as the ratio of displacement perpendicular to shear flow to that along shear flow. Our results showed that the directionality of macrophages was slightly but significantly enhanced by ox-LDL stimulation, implying that CD36^+^ macrophages exhibited a propensity to migrate in the vertical direction (Figure 3H). We also compared a key dynamic parameter of transmigration duration, which is defined as the interval from the timepoint of cell arrival at the site to the endpoint of completion of transmigration beneath the endothelial monolayer (Figure S6H). The result showed that transmigration duration was significantly shortened in CD36^+^ macrophage (Figure 3I).

Considering that protrusion is critical for cell crawling and migration, we then investigated whether intracellular lipid could affect the formation of protrusions. The number and the length of protrusions, and cell area were measured using the ImageJ software (Figure S6I). The results showed that the numbers, as well as the lengths of protrusions were significantly increased in CD36^+^ macrophages (Figure 3J and 3K). We further quantified macrophagic cell area and found that the mean area of CD36^+^ macrophages was significantly larger than the control (Figure 3L). Additionally, we examined various integrins and adhesive ligands expressed on the cell surface, which have been reported to be involved in adhesion and migration.^43^ The results showed that the expression levels of α-integrins, β-integrins and intercellular adhesion molecules were significantly upregulated in the infected CD36^+^ macrophages (Figure 3M and S6J). Additionally, these data were corroborated by the circulating CD36^+^ macrophages sorted from peripheral blood (Figure S6K). Further, we tested the adhesion and movement ability of CD36^+^ macrophages when adhesion molecule ICAM-1 was blocked. The data showed that the cellular adhesion ability, crawling displacement and crawling speed were significantly impaired, whereas the directionality and transmigration duration of macrophages were barely affected by ICAM1 antibody treatment (Figure S6L-S6S). Collectively, these data suggested that the foaminess of macrophages greatly enhanced their ability to adhere under mechanical stress, predisposing them to stay and to migrate vertically on the surface of BMECs; this might be attributable to the formation of protrusions as well as the expression of integrins and adhesion ligands.

### β-hydroxybutyrate promotes CD36^+^ macrophage formation and exacerbates neuroinvasion

Next, we performed lipid metabolomics analysis of serum samples to reveal how CD36^+^ macrophages were formed during bacterial neuroinvasion. Principal component analysis (PCA) showed that distinct metabolite genotype clusters were generated in the neurolisteriosis group (Figure 4A). The volcano plot indicated that 70 fatty acids (FAs) were upregulated and 71 FAs were downregulated (p < 0.05, |log_2_(fold change) | > 1) in the neurolisteriosis group (Figure 4B, Table S1). The 70 upregulated FAs were classified into four subclusters by the number of carbon atoms in the alkyl chain. The synthesis and metabolism cycle of long-chain FAs is longer than that of short-chain FAs. Based on the consideration of rapid progression of bacterial neuroinvasion, we focused on short-chain FAs and found that β-hydroxybutyrate (BHB), which has been reported to favor the foamy-like phenotype of macrophages ^44^, was the only short-chain FA among all upregulated FAs (Figure 4C). Kyoto Encyclopedia of Genes and Genomes (KEGG)-based analysis revealed that metabolites related to the synthesis and degradation of ketone bodies were enriched (Figure S7A), and metabolomics data showed that BHB was significantly increased during neuroinvasion (Figure S7B). Further, we found that BHB levels in mice serum climbed steadily over time in the acute neuroinvasion model of *L. monocytogenes* infection (Figure 4D). This finding is consistent with the increases in the bacterial load in the brain and the proportion of CD36^+^ macrophages in the PBMCs (Figure S7C and S7D).

**Figure 4.**
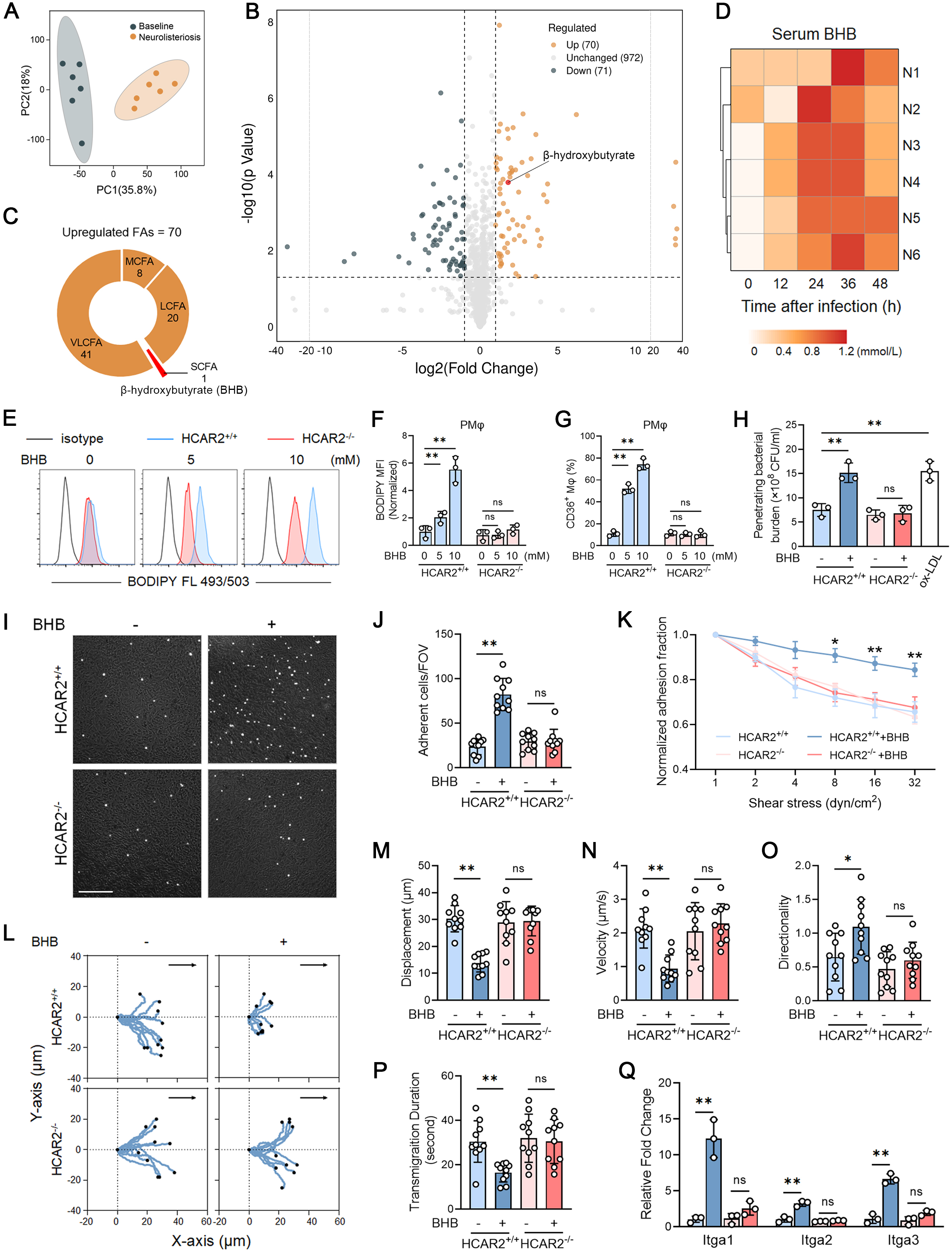
β-hydroxybutyrate aggravates bacterial neuroinvasion by promoting the formation of CD36^+^ macrophages via HCAR2 receptor. (A-C) Lipid metabolomic profiling analyses of serum from mice with or without neurolisteriosis (n=6 for each group). (A) Principal coordinates analysis (PCA) analysis of differences between groups. (B) Expression of FAs in serum was shown in the volcano plot. Differentially expressed FAs were selected with |log_2_ fold change|>1 and adjusted P < 0.05. The dark blue dots represent down-regulated FAs and the orange dots represent up-regulated FAs. (C) Classification of up-regulated FAs (SCFA, short chain FAs; MCFA, medium chain FAs; LCFA, long chain FAs; VLCFA, very long chain FAs) shown in the pie chart. (D) Biochemical analysis of serum BHB levels in mice during neurolisteriosis. (E-Q) Primary peritoneal macrophages were extracted from HCAR2^+/+^ and HCAR2^-/-^mice in preparation for further functional analysis. HCAR2^+/+^ and HCAR2^-/-^ macrophages were pretreated with BHB (10 mM) for 24 hours and then infected with *L. monocytogenes* (MOI=10) for 6 hours. (E-F) Flow cytometry analysis (E) of BODIPY FL and normalized MFI (F) in BHB-treated macrophages (0, 5, 10 mM). (G) Percentage of CD36^+^ macrophages in BHB-treated macrophages by flow cytometry. (H) Quantification of penetrating bacterial burden in the neuron cells in the lower chamber of Transwell system. The macrophages stimulated with ox-LDL (80 ng/ml) were used as positive control. (I-J) Optical images (I) of adherent macrophages on the surface of monolayer endothelial cells. Scale bar, 200 μm. Quantification of adherent cells in each field of view (J). (K) Macrophages resistance to shear flow after adhesion to endothelial cells. (L-P) Crawling and migrating dynamics of infected macrophages under shear flow. (L) Typical trajectories of infected macrophages crawling on monolayer endothelial cells. Quantification of crawling displacement (M), velocity (N), directionality (O) and transmigration duration (P). (Q) MRNA expressions of intercellular α-integrin in infected macrophages. Experiments were performed in duplicate (D) or triplicate (E-Q). Data are presented as mean ± SEM [n=3 in (F-H, Q), n=10 in (J, M-P)]. *p < 0.05, **p < 0.01. Abbreviations: ns, nonsignificant; FAs, fatty acids; BHB, β-hydroxybutyrate; MFI, mean value of fluorescence intensity.

Subsequently, we sought to explore the reasons for the enhanced ketogenesis during bacterial brain invasion. Liver tissues from acute neuroinvasion mice model were collected for transcriptome analysis by RNA-seq. The results showed that compared with controls, the genes associated with ketogenesis were sharply upregulated in the livers of mice suffered from neurolisteriosis, whereas these genes were moderately increased in the mildly infected group (Figure S7E). KEGG-based analysis and gene-set enrichment analysis (GSEA) between the neurolisteriosis and control group revealed that the differentially expressed (DE) genes were enriched in glycolysis or gluconeogenesis (Figure S7F and S7G). To validate this finding, we measured serum glucose levels in mice and found that glucose levels decreased significantly and remained low during *L. monocytogene*-induced neuroinvasion (Figure S7H); meanwhile, we observed a significant decrease in cellular energy and an increase in ADP/ATP ratios in mice with neurolisteriosis compared to controls (Figure S7I and S7J). These data implied that the decrease in glucose levels and increase in ketogenesis may be induced by excessive glycolysis during *L. monocytogene*-induced neuroinvasion. Further, we showed that the extracellular acidification rates (ECAR) were significantly increased in the livers of mice with neurolisteriosis (Figure S7K). This data was confirmed by the measurement of lactate, the end product of glycolysis (Figure S7L). Also, we found that mice experienced weight loss and reduced food intake, which to a certain extent contributed to ketogenesis after infection (Figure S7M and S7N). Interestingly, there was no significant difference between the mildly infected group and the control group in these indicators during the experiments (Figure S7H-S7N). Furthermore, a steady increase in serum BHB and a decrease in glucose levels were also observed in the neuroinvasion models established using *S. typhimurium* and MRSA (Figure S7O and S7P). These data indicated that BHB production was significantly induced during bacterial neuroinvasion, possibly due to low serum glucose levels induced by excessive glycolysis.

As previously reported, autocrine or paracrine BHB directly activates macrophages function mainly via the hydroxycarboxylic acid receptor 2 (HCAR2).^45^ Therefore, we next used primary peritoneal macrophages isolated from HCAR2 knockout mice to study whether BHB could promote bacterial neuroinvasion. We confirmed that lipid accumulation and percentage of CD36^+^ macrophages all increased dramatically upon BHB stimulation, and there was no significant difference when HCAR2 was knocked out (Figure 4E-4G, S7Q). BHB inhibits adenylyl cyclase activity through the HCAR2 receptor and reduces cellular cyclic adenosine monophosphate (cAMP) levels, thereby protecting lipid droplets from lipolysis by hormone-sensitive lipases (HSL).^45^ As a result, we found that cAMP pretreatment eliminated the decrease in HSL activity, the increase in lipid content and intracellular bacterial burden stimulated by BHB; however, this phenomenon was not found in HCAR2-deficient macrophages (Figure S7R-S7T). By using the *in vitro* BBB model (Figure 2A), the number of bacteria crossing the barrier increased significantly in response to BHB stimulation in controls, but not in HCAR2^-/-^ macrophages (Figure 4H). Next, peritoneal macrophages from HCAR2^+/+^ and HCAR2^-/-^ mice were isolated to further assess the effects of BHB on the biophysical dynamics of macrophages. The data demonstrated that BHB significantly altered the static or shear-resistant adhesion of control macrophages, but not of HCAR2^-/-^ macrophages (Figure 4I-4K). We also found that BHB greatly impaired the crawling ability, including shortened crawling displacement, slowed down crawling speed and unpredictable directionality of macrophages (Figure 4L-4O). In addition, the transmigratory ability of macrophages was significantly increased when cells were pretreated with BHB (Figure 4P). Consistently, the expression levels of the α-integrin family were significantly upregulated by BHB treatment in controls, but not in HCAR2-deficient macrophages (Figure 4Q). Taken together, these data indicated that BHB facilitated the formation of CD36^+^ macrophages through the HCAR2 receptor, thereby affecting the process of bacterial neuroinvasion.

### BHB promotes the survival of macrophages via HCAR2-mediated calcium influx

Because the foamy stage appears to be a terminal point of macrophage differentiation ^46^, we wondered whether, in addition to converting peritoneal macrophages to the foamy phenotype, cell survival also contributed to the increase in absolute quantity of CD36^+^ macrophages induced by BHB treatment. Immunofluorescence was performed with the objective of showing the presence of *L. monocytogenes* within the macrophages (Figure S8A). The results showed that the lactate dehydrogenase (LDH) activity in the culture medium was significantly reduced in response to BHB treatment, indicating a decrease in cell death that induced by infection (Figure 5A).

**Figure 5.**
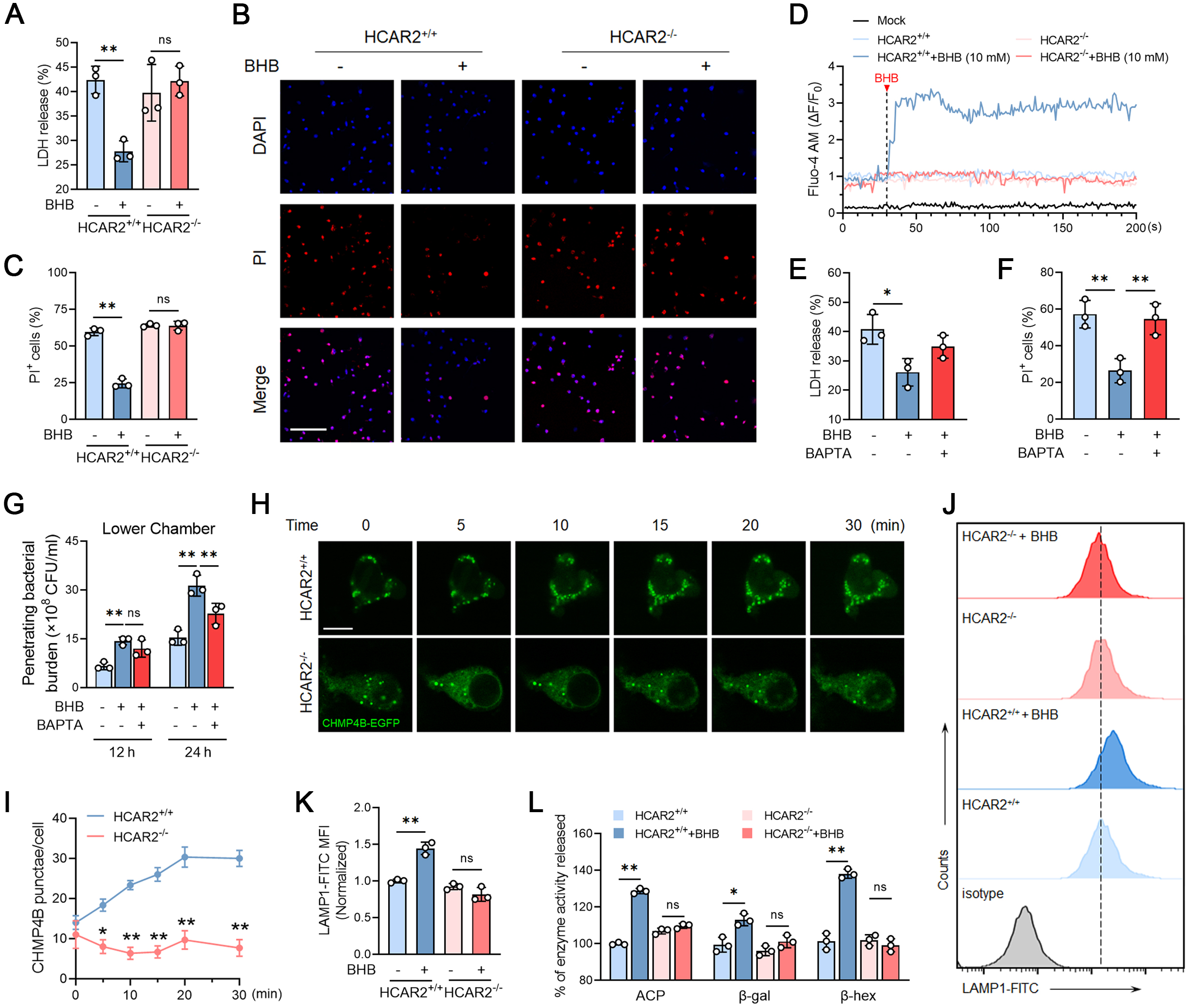
β-hydroxybutyrate promotes the survival of macrophages via HCAR2-mediated calcium influx. (A-L) Primary peritoneal macrophages were extracted from HCAR2^+/+^ and HCAR2^-/-^ mice. Cells were then infected with *L. monocytogenes* (MOI=10) for 6 hours. (A-C) HCAR2^+/+^ and HCAR2^-/-^ macrophages were pretreated with BHB (10 mM) before infection. (A) Quantification of the released LDH in cell culture medium. (B) Immunofluorescence of propidium iodide (PI)-stained macrophages. Scale bar, 100 μm. (C) Quantification of the PI^+^ cells after infection. (D) Flow cytometry analysis of dynamics in intracellular calcium concentrations. Infected macrophages were preloaded with Fluo-4 AM. BHB (10 mM) was added to the buffer 30 seconds later. (E-G) Macrophages were pretreated with BHB (10 mM) or BAPTA (10 μM) 24 hours before infection. (E) Quantification of the released LDH in cell culture medium. (F) Quantification of the PI^+^ cells post infection. (G) Quantification of penetrating bacterial burden in the neuron cells in the lower chamber of Transwell system. (H-I) Infected macrophages were treated with BHB (10 mM). (H) Time-lapse images of CHMP4B-EGFP in macrophages were collected every 5 minutes post BHB treatment with confocal microscopy. (I) Quantification of CHMP4B-EGFP puncta. (J-L) HCAR2^+/+^ and HCAR2^-/-^ macrophages were pretreated with BHB (10 mM) for 24 hours before infection. Flow cytometry analysis (J) and the normalized MFI (K) of LAMP1 expression on the cell surface. (L) Quantification of the lysosomal enzyme [acid phosphatase (ACP), β-galactosidase (β-gal) and β-hexosaminidase (β-hex)] activities released into the culture medium. Experiments were performed in triplicate. Data are presented as mean ± SEM (n=3). *p < 0.05, **p < 0.01. Abbreviations: ns, nonsignificant; BHB, β-hydroxybutyrate; LDH, lactate dehydrogenase.

Propidium iodide (PI) staining was used to evaluate the membrane integrity of infected macrophages, and we found that the proportion of PI^+^ cells decreased from 59.7% to 24.5% under BHB stimulation, with no significant difference in cell viability between the BHB treatment and control groups when HCAR2 was knocked out (Figure 5B and 5C). Calcium influx is generally recognized as a signal that triggers cell membrane repair.^47^ To investigate the impact of BHB-HCAR2 signaling on cytosolic calcium concentration, infected macrophages were loaded with Fluo-4 AM, a fluorescent Ca^2+^ indicator. The results showed that BHB induced remarkable accumulation of intracellular calcium, and HCAR2 knockdown eliminated calcium influx in macrophages (Figure 5D). Pretreatment of macrophages with BAPTA, a Ca^2+^ chelator, reversed the BHB-induced increase in cell survival (Figure 5E and 5F). Further, we tested whether calcium flux was associated with CD36^+^ macrophage-mediated intracellular bacterial penetration of the BBB. As a result, inhibition of calcium flux had no effect on bacterial uptake by macrophages in the upper chamber (Figure S8B); however, we observed differences in bacterial loads in the macrophages that had crossed the barrier (Figure 5G).

We next investigated how BHB maintained the survival of CD36^+^ macrophages. As previously reported, calcium influx initiates membrane repair mainly through recruitment of the endosomal sorting complexes required for transport (ESCRT) machinery.^48^ Our data showed that BHB treatment significantly increased the number of puncta that were positive for charged multivesicular body protein 4B (CHMP4B), which indicated assembly of ESCRT in the infected macrophages; HCAR2 knockdown or BAPTA treatment could block this membrane repair process (Figure 5H and 5I, S8C and S8D). Cellular membrane repair also relies on lysosomes, which can rapidly fuse with the plasma membrane when intracellular Ca^2+^ concentration rises. Therefore, we examined lysosomal exocytosis in the infected macrophages using exofacial lysosomal-associated membrane protein 1 (LAMP1) and the release of lysosomal enzymes, such as acid phosphatase (ACP), β-galactosidase (β-gal) and β-hexosaminidase (β-hex). The results confirmed that BHB promoted the lysosomal exocytosis in infected macrophages through the HCAR2 receptor (Figure 5J-5L, S8E-S8G). As thioglycolate has been demonstrated to initiate an inflammatory response and alter lysosomal enzymes during the induction of PBMCs into macrophages ^49^, we further detected the lysosomal exocytosis of infected resident peritoneal macrophages and found that BHB indeed promoted the lysosomal exocytosis, thereby maintaining the survival of resident CD36^+^ macrophages (Figure S8H-S8J). Lipophagy has been reported to implicate in regulating lipid metabolism and cell survival, particularly during intracellular bacterial infections ^50^, we asked whether lipophagy was involved in modulating phagocyte survival during infection. The results showed that lipophagy was significantly activated in CD36^+^ macrophages following BHB treatment.

However, this activation was abrogated in the HACR2 knockout group (Figure S8K). Importantly, we found that the proportion of CD36^+^PI^+^ macrophages was reduced after BHB treatment, and that the effect of BHB was significantly inhibited by pretreatment with 100 nM lipophagy inhibitor Baf-A1 for 6 hours (Figure S8L). The results suggested that BHB promotes macrophage survival, which is likely to be dependent on lipophagy. Collectively, these data indicated that BHB signaling activated Ca^2+^-mediated membrane repair, thereby maintaining the survival of CD36^+^ macrophages via the HCAR2 receptor.

### Ketogenesis exacerbates intracellular bacterial neuroinvasion

BHB can be synthesized via the metabolism of FAs and ketogenesis.^51^ To elucidate the correlation between ketogenesis and bacterial neuroinvasion, mice were administered varying doses of a ketone ester drink for a period of two weeks, after which they were infected with 2×10^8^ CFU of *L. monocytogenes* via intraperitoneal injection (Figure 6A). The data showed that the ketone ester elevated the levels of serum BHB and the proportion of CD36^+^ macrophages in peritoneal blood (Figure 6B and 6C), accompanied by remarkable increases in the clinical scores, mortality rate and bacterial load in brain tissues (Figure S9A, 6D and 6E). It has been established that dietary modulation can increase and maintain circulating metabolites in the human circulation, and a growing body of evidence demonstrates that a ketogenic diet elevates BHB levels.^52^ The ketogenic diet is a low-carbohydrate, high-fat diet that has been demonstrated to result in weight loss and confer a multitude of health benefits, including the alleviation of symptoms associated with neurodegenerative diseases, cardiovascular diseases, and cancers.^21,53–55^ We proceeded to investigate whether a ketogenic diet was associated with neurolisteriosis. The HCAR2^-/-^ mice and their littermates were fed a normal or ketogenic diet for four weeks, respectively, and then infected with *L. monocytogenes* via intravenous injection (Figure 6F). Consequently, the lifespan of HCAR2^+/+^ mice fed a ketogenic diet was markedly reduced in the presence of *L. monocytogenes* infection, and the detrimental impact of dietary modulation was abrogated by HCAR2 knockout (Figure 6G). Consistently, the clinical scores (Figure S9B), percentages of CD36^+^ macrophages in peritoneal blood (Figure 6H), the intracellular bacterial burden of macrophages (Figure 6I), and bacterial loads in brain tissues (Figure 6J) were all significantly increased in HCAR2^+/+^ mice on a ketogenic diet. Interestingly, mice in this group exhibited less weight loss than mice in the control group, and experienced greater dissemination of bacteria to other organs (Figure 6J and S9C). Subsequently, the levels of pro-inflammatory factors in the CSF and brain homogenate were examined. The results showed that a ketogenic diet significantly increased the levels of pro-inflammatory cytokines and precipitated more pronounced neuronal injury in HCAR2^+/+^ mice, whereas it had no discernible impact on HCAR2^-/-^ mice (Figure 6K-6N). Notably, no extravasation of Evans blue was observed in the brains of mice fed a ketogenic diet, thereby ruling out the possibility of direct damage to the BBB (Figure 6M and 6O). These data were corroborated by an intraperitoneal injection of 2×10^8^ CFU of *L. monocytogenes* in the same experimental model (Figure S9D-S9G). Further, the inflammatory profile of the CD36^+^ macrophages generated from mice fed a ketogenic diet was examined. It was observed that these CD36^+^ macrophages also exhibited an anti-inflammatory phenotype (Figure S9H), and the inflammatory signature was not affected by BHB-HCAR2 signaling once CD36^+^ macrophages was formed (Figure S9I and S9J). These characteristics were identical to those of CD36^+^ macrophages isolated from mice following infection (Figure S2M, S3F and S3G).

**Figure 6.**
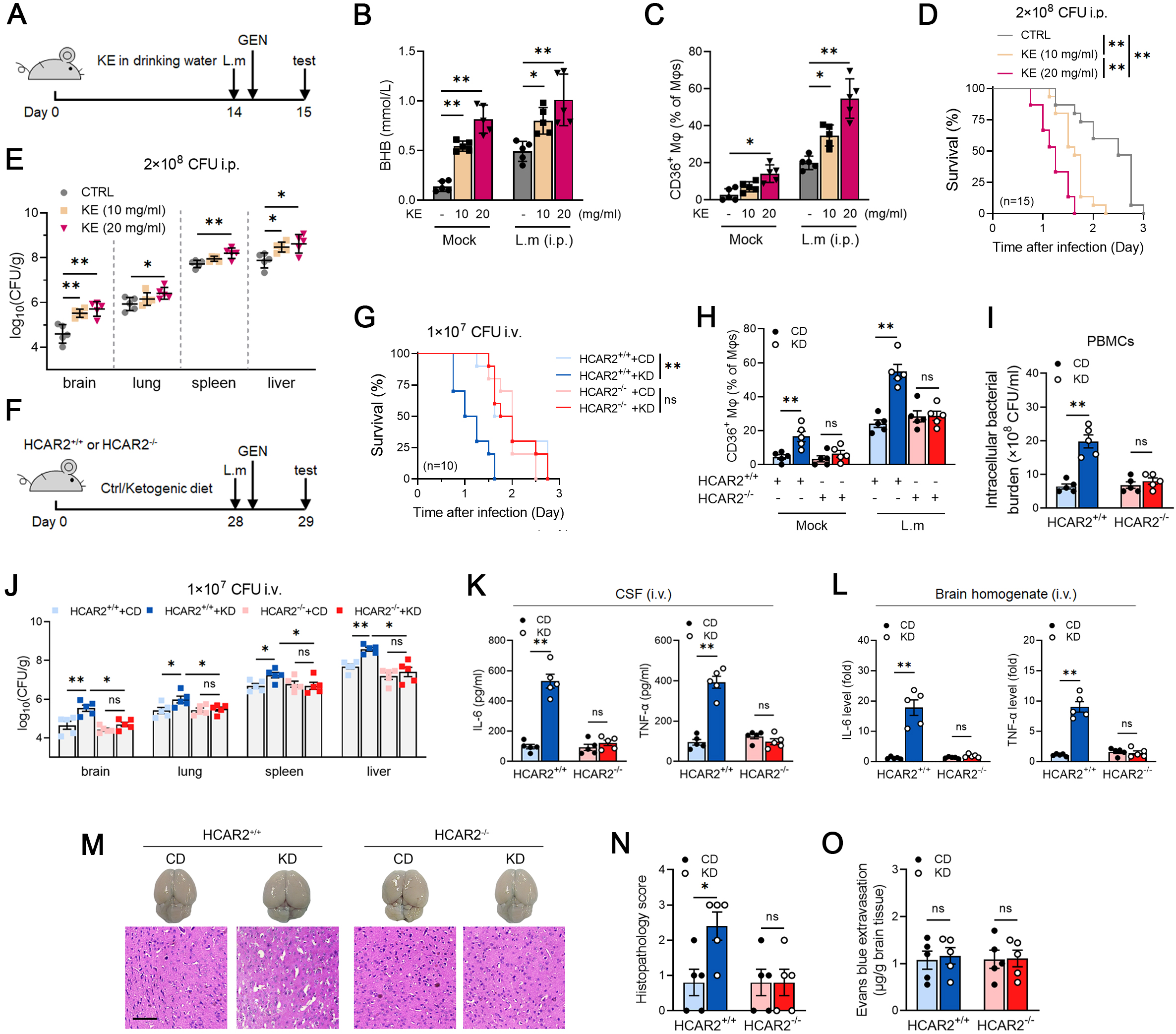
Ketogenesis exacerbates neurolisteriosis through β-hydroxybutyrate-HCAR2 signaling. (A-D) (A) Schematic diagram of KE supplementation in mice. C57BL/6 mice were supplemented with KE (10, 20 mg/ml) in drinking water for 2 weeks and then intraperitoneally challenged with 2×10^8^ CFU *L. monocytogenes*. Gentamicin (30 mg/kg) was administered intraperitoneally 4 hours after infection. (n=15 for each group) (B) Serum BHB levels. (C) Percentage of CD36^+^ macrophages. (D) Survival of mice exposed to *L. monocytogenes*. (E) Bacterial burdens in the organs. (F-L) (F) Schematic diagram of ketogenic diet in mice. HCAR2^+/+^ and HCAR2^-/-^ mice were treated with ketogenic or control diet for 4 weeks, followed by intravenous infection of 1×10^7^ CFU *L. monocytogenes*. Gentamicin (30 mg/kg) was administered intraperitoneally 4 hours after infection. Samples were collected for examination 24 hours post infection. (n=10 for each group) (G) Survival of mice exposed to *L. monocytogenes*. (H) Percentage of CD36^+^ macrophages by flow cytometry. (I-J) Bacterial burdens in PBMCs (I) and parenchymal organs (J). (K-L) Pro-inflammatory factor levels in CSF (K) and brain homogenates (L). (M-O) Representative images (M) of Evans blue leakage in mouse brains and HE staining in cerebral cortex tissues. Scale bar, 10 μm. Quantification of pathological score (N) and Evans blue leakage (O). Experiments were performed in duplicate. Data are presented as mean ± SEM (n=5). *p < 0.05, **p < 0.01. Abbreviations: ns, nonsignificant; KE, Ketone ester; GEN, gentamicin; CD, control diet; KD, ketogenic diet; i.p., intraperitoneally; i.v., intravenously.

Further, to exclude the possibility of BHB exerting its effects by preventing the clearance of bacteria before they reach the brain, we isolated peritoneal macrophages from HCAR2^+/+^ and HCAR2^-/-^ mice to perform adoptive macrophage transfer experiments (Figure S9K). Gentamicin was used to eliminate circulating bacteria that had not yet entered the phagocytes. The results showed that mice transferred with macrophages obtained from HCAR2^+/+^ mice fed a ketogenic diet had a shortened survival time in comparison to mice fed a control diet. However, this difference in survival time between different diets was eliminated in the HCAR2^-/-^ group (Figure S9L). The observation was further confirmed by examining bacterial loads in the brain tissues, and inflammatory factor levels in the CSF of recipient mice (Figure S9M and S9N). In addition to HCAR2, BHB has been shown to act through other mechanisms, therefore a 3-hydroxy-3-methylglutaryl-CoA synthase 2 (HMGCS2, the rate-limiting enzyme during ketogenesis) knockout mouse line was used to validate our findings (Figure S9O). The results showed that intravenous infection with 1×10^7^ CFU of *L. monocytogenes* resulted in a dramatic decrease in blood BHB levels and the proportion of CD36^+^ macrophages in the peritoneal blood of the Hmgcs2 knockout mice compared with the controls (Figure S9P and S9Q), accompanied by a significant reduction in mortality rates, brain tissue bacterial loads, and CSF inflammatory factor levels (Figure S9R-S9T). These data suggested that BHB may exert its effects though promoting the formation of CD36^+^ macrophages via HCAR2 receptor, thus exacerbating bacterial neuroinvasion. We also employed MRSA to further verify our findings. The data indicated that, as expected, the ketogenic diet facilitated brain invasion by MRSA and intensified the phenotypes of neuroinvasion (Figure S9U-S9W). Taken together, our data demonstrated that a ketogenic diet could exacerbate bacterial neuroinvasion and hasten the demise of mice, primarily depending on the elevation in serum BHB via the HCAR2 receptor.

Our data thus far suggest that BHB plays an important role in bacterial neuroinvasion in mice, and that the enhancement in ketogenesis is probably induced by low serum glucose levels. Subsequently, the impact of restoring blood glucose levels on the outcomes of infected mice was evaluated. The results showed that in an acute neuroinvasion model of *L. monocytogenes* infection, the supplementation of mice with glucose at 12-hour intervals successfully restored blood glucose levels and inhibited hepatic glycolysis (Figure S10A-S10F), leading to significant decreases in BHB levels and CD36^+^ macrophage percentages (Figure S10G-S10H). Surprisingly, we found that four rounds of glucose supplementation significantly increased the survival of mice, and reduced their clinical scores, brain bacterial burden and pro-inflammatory cytokines levels (Figure S10I-S10M). These data indicate that glucose homeostasis may limit the brain invasion by *L. monocytogenes* infection and alleviate neuroinvasion symptoms in mice.

### Serum BHB fuels the symptoms of human bacterial neuroinvasion

A retrospective study based on the Medical Information Mart for Intensive Care IV (MIMIC-IV) was performed to explore the risk factors for bacterial neuroinvasion in humans. As a result, ferritin (OR 5.09, 95% CI 1.55-16.71), fibrinogen (OR 3.57, 95% CI 1.13-11.25), LDL (OR 3.31, 95% CI 1.01-10.83), glucose (OR 2.17, 95% CI 1.04-4.50) and aspartate amino transferase (OR 2.16, 95% CI 1.01-4.63) were identified as independent risk factors, whereas potassium (OR 0.42, 95% CI 0.20-0.87), alkaline phosphatase (AP, OR 0.38, 95% CI 0.18-0.83), phosphate (OR 0.23, 95% CI 0.11-0.50), high-density lipoprotein (HDL, OR 0.19, 95% CI 0.05-0.66) and transferrin (OR 0.14, 95% CI 0.04-0.50) were identified as protective factors in patients with bacterial neuroinvasion (Figure S11A). Next, we asked whether the accumulation of BHB or CD36^+^ macrophages could also be identified and were associated with bacterial infection of the CNS in human clinical samples. Gender- and age-matched bacterial neuroinvasion subjects and healthy individuals were recruited and peripheral blood samples were collected for analysis (Figure 7A). Since blood glucose is a risk factor for bacterial neuroinvasion and BHB level can be dramatically affected by glycolysis, subjects with abnormal blood glucose or insulin concentrations were excluded from the two groups (Figure S11B-S11E). As a result, serum BHB levels were significantly elevated in patients compared with healthy controls, suggesting an infection-induced ketogenesis (Figure 7B). Interestingly, serum cholesterol levels, including total cholesterol (TC) and LDL-C were increased in patients compared with healthy controls, while HDL-C was slightly but not significant reduced (Figure S11F-11H). This finding is consistent with data from our retrospective study based on MIMIC-IV (Figure S11A). To investigate whether BHB was associated the severity of infection, we further divided patients into three groups, high BHB (in the top 1/3 among samples tested, n=21), moderate (middle 1/3, n=20) and low (bottom 1/3, n=21; Figure 7A). Clearly, patients with high BHB levels exhibited higher clinical scores than patients with low BHB levels (Figure 7C). Consistently, high-BHB patients had higher cell counts and more protein in the CSF (Figure 7D and 7E); higher infection indices such as white blood cell counts (WBC; Figure 7F) and procalcitonin (PCT; Figure 7G); and higher values for markers of inflammatory conditions such as erythrocyte sedimentation rate (ESR; Figure 7H) and C-reactive protein (CRP; Figure 7I), suggesting increased disease severity. These data implied that infection-induced ketogenesis might provide important fuel for severe bacterial neuroinvasion.

**Figure 7.**
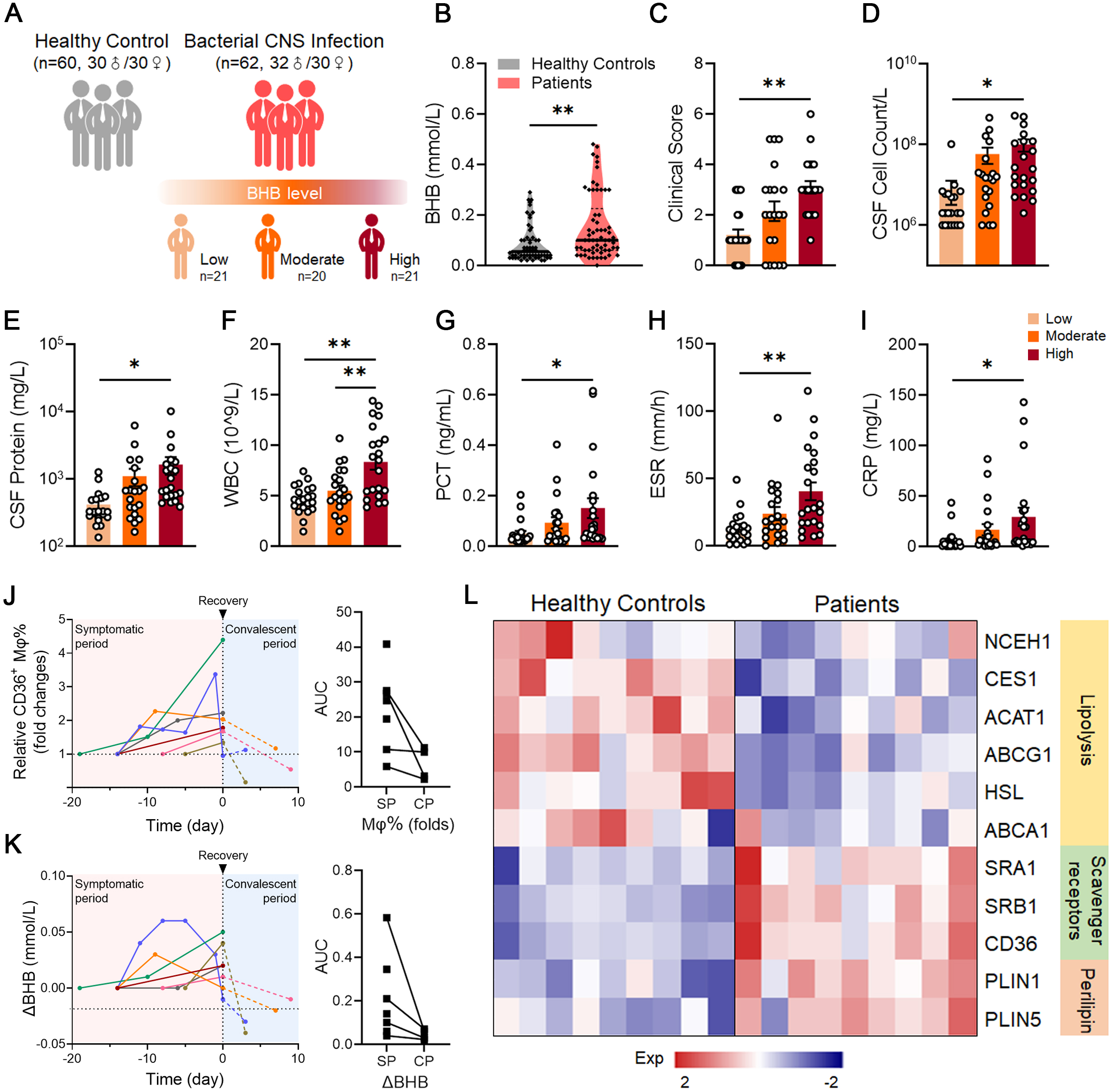
β-hydroxybutyrate levels are increased during bacterial neuroinvasion in humans. (A) Schematic diagram of samples from healthy controls (n=60) and patients with bacterial CNS infection (n=62). The patients were divided into three groups according to serum BHB levels. (B) Serum BHB levels in healthy controls and patients. (C-I) Clinical scores (C), cell counts (D) and protein levels (E) in CSF. White blood cells (WBC) (F), procalcitonin (PCT) (G), erythrocyte sedimentation rate (ESR) (H) and C-reactive protein (CRP) (I) in the peripheral blood of patients with different BHB levels. (J-K) Dynamics of CD36^+^ macrophages (J) and serum BHB (K) in 7 donors after admission. The time after admission was divided into symptomatic and convalescent periods according to clinical and laboratory indexes of infection. The scatter plot on the right panel represents the statistics of AUC. (L) Relative expression of genes involved in lipid metabolism in PBMCs from healthy controls and patients was detected by qRT-PCR. Data are presented as mean ± SEM. *p < 0.05, **p < 0.01. Abbreviations: ns, nonsignificant; AUC, area under the curve; BHB, β-hydroxybutyrate.

To explore the dynamics of CD36^+^ macrophages and BHB in the pathogenesis of bacterial neuroinvasion, we followed-up with 7 individuals during their hospitalization. We found that CD36^+^ macrophages and BHB levels in patients tended to increase during disease progression, and decrease during the recovery stage (Figure 7J and 7K). The intracellular bacterial load of CD36^+^ macrophages sorted from patients was also calculated. The results showed that bacteria were predominantly present in CD36^+^ macrophages, but not in CD36^-^ macrophages (Figure S11I). To further confirm the relationship between macrophage lipid metabolism and bacterial neuroinvasion pathology, we collected PBMCs from patients or healthy controls for analysis. Compared with healthy controls, the expression levels of genes involved in cholesterol uptake were significantly increased, while those of genes related to cholesterol utilization and excretion were decreased in macrophages from patients (Figure 7L). Taken together, these data indicated that BHB fueled human bacterial neuroinvasion by promoting the development of CD36^+^ macrophages.

## Discussion

Our findings suggested that CD36^+^ macrophages may serve as a carrier for intracellular bacteria that colonize the brain, thereby facilitating the penetration of these bacteria across the BBB. Generally, the BBB prevents pathogens from crossing the brain tissue unless there is significant trauma to the brain. Nevertheless, specific bacteria are capable of breaking through the BBB via their distinctive mechanisms, leading to severe brain infections.^15,56^. In this study, we discovered that intracellular bacteria may prefer to exploit a “Trojan horse” pathway, which causes minimal damage to the BBB but enables acute brain invasion. It is possible that CD36^+^ macrophages may be hijacked by intracellular bacteria and serve as vehicles for neuroinvasion. In comparison to peritoneal macrophages, CD36^+^ macrophages exhibit a greater number of protrusions and a larger cell area, which renders them more conducive to colonization on BMECs under the pressure of blood flow (Figure 3E-3L). Based on the aforementioned remarkable characteristics, CD36^+^ macrophages have the potential to become carriers of engineered intracellular bacteria for bacteriotherapy.

Pathogens have the capacity to exploit the majority of the host’s nutrients, including glycerol, FAs, and a number of other carbon-based substances, with the objective of reprogramming immune cells.^57^ Our findings have demonstrated that intracellular bacteria are capable to actively influence host free FA metabolism (Figure 4A-4C) and reprogram macrophages, thereby exacerbating neuroinvasion. The foamy-like phenotype of macrophages is formed through dysregulation of lipid metabolism, which in turn impairs the immune function of macrophages and contributes to pathogenesis. For example, foamy macrophage necrosis has previously been observed to expand tuberculous lesions in lung parenchyma, resulting in progressive destruction of lung tissue and loss of lung function in patients with active disease.^58^ In this study, we discovered that BHB produced by a ketogenic diet promoted the formation of a foamy-like phenotype of macrophages in peripheral blood, thereby facilitating bacterial neuroinvasion. Interestingly, we found that serum cholesterol levels, including TC and LDL-C were significantly higher in patients than in healthy controls (Figure S11F and S11H). We hypothesize that cholesterol and other lipid metabolites may also be involved in the formation of CD36^+^ macrophages. However, this hypothesis requires further experimental verification.

The biomechanical data indicated that CD36^+^ macrophages exhibited a greater degree of adhesion to the endothelial surface than normal macrophages, and this adhesion-enhancing effect was more pronounced when endothelial cells were dense (Figure 3B and 3C, S6B and S6C). Accordingly, we found that an elevation in CD36^+^ macrophages in peripheral blood resulted in the spread of bacteria predominantly to the brain, as opposed to other organs (Figure 2G and S3Q). This suggested that CD36^+^ macrophage-mediated bacterial invasion affected the brain to a greater extent than other tissues. Interestingly, it was observed that a ketogenic diet and ketone ester supplementation significantly increased bacterial load not only within the brain, but also in other tissues such as the liver and lung (Figure 6E and 6J). This suggested that, in addition to promoting the generation of CD36^+^ macrophages, ketogenesis might play a role in the intracellular bacterial infections that is currently unknown. Further studies are therefore necessary to determine the role of ketogenesis in the formation of CD36^+^ macrophages and the contribution of ketogenesis to bacterial neuroinvasion.

The basal nutritional status of the host may also have an important influence on the outcome of infection. In this study, we showed that neurolisteriosis mice experienced a decrease in blood glucose levels and an increase of ketogenesis in the liver (Figure S7A-S7H). We also found that during the course of bacterial neuroinvasion, mice exhibited a reduction in body weight and diminished appetite (Figure S7M and S7N). Interestingly, the fluctuations in glucose levels appeared to precede a decline in food intake in neurolisteriosis mice (Figure S7H and S7N). In this context, the decrease in glucose levels and increase in ketogenesis cannot be attributed exclusively to anorexia, and further studies are therefore required to elucidate the underlying mechanisms of the fluctuations in glycolysis and glucose levels during neuroinvasion.

Our findings suggest that the maintenance of optimal blood sugar levels may be a key factor in the onset, progression, and prognosis of bacterial neuroinvasion. Several recent reports provide support for this hypothesis. For example, glucose supplementation has been shown to exert a protective effect against *Yersinia pseudotuberculosis* infection in mice.^59^ In addition, reduced BHB levels in patients with sepsis have been demonstrated to diminish glucose production, so that the blood glucose levels in non-survivor sepsis patients are likely to be considerably lower than in the survivors. ^60–62^ Interestingly, another study by Wang et al. showed that glucose supplementation was detrimental in bacterial sepsis. However, in their study, there was no discernible difference in the kinetics of blood glucose following glucose treatment ^63^, which further suggests the pivotal role of blood glucose levels in bacterial infections. Consequently, we endeavored to replenish blood sugar levels in mice afflicted with acute neuroinvasion via intraperitoneal administration of glucose, and surprisingly found that glucose supplementation resulted in a marked alleviation of the clinical symptoms, a reduction in the bacterial burden within the brain, and an improvement in the survival rate of the mice (Figure S10). These findings highlight the crucial role of nutritional therapy, in addition to antibacterial therapy, in the clinical treatment of bacterial neuroinvasion. In addition, *L. monocytogenes* has been shown to induce the accumulation of lipid droplets in macrophages, thereby contributing to bacterial survival and evasion of innate immunity. ^64^ Given that intracellular lipid is the key regulator of CD36^+^ macrophage function, targeting this lipid utilization pathway may be a promising therapeutic strategy for bacterial neuroinvasion.

Overall, this study demonstrated that a BHB-rich environment, as created by the ketogenic diet, rendered mice more susceptible to *L. monocytogene*-induced neuroinvasion. From a mechanistic perspective, BHB protected CD36^+^ macrophages from the onset of senescence via HCAR2-mediated membrane repair. However, the inhibition of HCAR2 signaling or calcium influx was observed to reverse the improvement in CD36^+^ macrophage survival, yet did not fully reverse the exacerbation in bacterial neuroinvasion induced by BHB (Figure 5E-5G). These data suggested that bacterial brain invasion induced by BHB is not solely dependent on HCAR2 signaling, and that other potential mechanisms may also be involved in this process. Nevertheless, this hypothesis requires further experimental confirmation.

### Limitations of the study

The role of other cell clusters or FAs in bacterial neuroinvasion, beyond CD36^+^ macrophages and BHB, remains to be elucidated. To circumvent the potential for BHB to exert its effects through alternative target cell populations and organs, the utilization of a HCAR2 conditional knockout mouse line is imperative. Moreover, additional human neuroinvasion samples induced by intracellular bacteria other than MTB should be further collected and analyzed.

## Acknowledgments

This work was supported by the National Natural Science Foundation of China (No. 92369115, 82422048 and 92469110), the National Key Research and Development Program of China (No. 2024YFC2310800 and 2021YFC2301405), the Fundamental Research Funds for the Central Universities (No. 2024CDJXY-016), Natural Science Foundation of Chongqing, China (No. CSTB2023NSCQMSX0402), and Chongqing Talents: Exceptional Young Talents Project (No. cstc2021ycjh-bgzxm0099). The funders had no role in study design, data collection and analysis, decision to publish, or preparation of the manuscript.

## Author Contributions

H.W., Q.Z., X.L. and Z.S. conceived and designed the study. Z.S., H.Y., H.F., Q.H., B.F., Y.X. and D.G. performed the animal experiments. Z.S., K.Y., Q.H., Q.L., Y.D. and Xushuo Z. carried out healthy volunteer and patient recruitment and sample collection. Z.S., N.L. and X.S. carried out the live-cell flow chamber assays. H.W., Z.S., K.Y., H.Y. and Xiaokai Z. collected and analyzed the data. X.T., S.F., J. L., R.Y., W.G., X.M. and M.L. contributed to experimental material and insightful suggestions. H.W., X.L., Xushuo Z. and Z.S. wrote the manuscript. H.W. and Q.Z. coordinated the project. All authors read and approved the final manuscript.

## Materials and methods

## KEY RESOURCES TABLE

| REAGENT or RESOURCE | SOURCE | IDENTIFIER |
| --- | --- | --- |
| Antibodies |  |  |
| Anti-Human/Mouse CD11b, FITC | Multi Sciences | Cat# F11011B01 |
| Anti-Mouse F4/80, PE | Multi Sciences | Cat# F21480A02 |
| Anti-Mouse CD36, APC | eBioscience | Cat# 17-0362-82 |
| Anti-Mouse ITGA2, FITC | eBioscience | Cat# 11-5971-82 |
| Anti-Mouse ITGB1, FITC | eBioscience | Cat# 11-0291-82 |
| Anti-Mouse ICAM1, PE | eBioscience | Cat# 11-0541-82 |
| Anti-Mouse CD107a, FITC | Proteintech | Cat# 65050-1-Ig |
| Anti-Mouse CD36 | CST | Cat# 28109 |
| Anti-Mouse CD31 | CST | Cat# 15585 |
| Anti-Mouse PPARG | CST | Cat# 2435 |
| Anti-Mouse β-Actin | CST | Cat# 4970 |
| Anti-Mouse IgG, HRP | CST | Cat# 7076 |
| Anti-rabbit IgG, HRP | CST | Cat# 7074 |
| Anti-Listeria monocytogenes | This paper | N/A |
| Anti-Mouse FABP4 | Abcam | Cat# ab92501 |
| Anti-Mouse PCSK9 | Abcam | Cat# ab185194 |
| Anti- Mouse CD11b | Abcam | Cat# ab8878 |
| Anti-Mouse LC3B | Abcam | Cat# ab192890 |
| Anti-ICAM1 | Abcam | Cat# ab ab222736 |
| Anti-Mouse α-TUBULIN | Santa Cruz | Cat# sc-398103 |
| Bacterial strains |  |  |
| Listeria monocytogenes | ATCC | Cat# 19115 |
| MRSA USA300 | ATCC | Cat# TCH1516 |
| Salmonella Typhimurium | ATCC | Cat# SL1438 |
| Streptococcus pneumoniae | ATCC | Cat# 49619 |
| Chemicals and Reagents |  |  |
| Thioglycollate medium | Sigma-Aldrich | Cat# 9032 |
| Collagenase IV | Sigma-Aldrich | Cat# C4-BIOC |
| Oxidized Low Density | Invitrogen | Cat# L34357 |
| Lipoprotein (ox-LDL) |  |  |
| DiI-OxLDL | Invitrogen | Cat# L34358 |
| DNase I | Invitrogen | Cat# 18047019 |
| Hoechst 34580 | Invitrogen | Cat# H21486 |
| Propidium iodide | Invitrogen | Cat# P1304MP |
| Fluo-4 AM | Invitrogen | Cat# F14217 |
| Lipofectamine 2000<br>Transfection Reagent | Invitrogen | Cat# 11668030 |
| Dulbecco's modified Eagle's<br>medium (DMEM) | GIBCO | Cat# C11965500BT |
| Fetal bovine serum (FBS) | GIBCO | Cat# 10091148 |
| Penicillin Streptomycin (Pen<br>Strep) | GIBCO | Cat# 15140122 |
| Collagenase II | GIBCO | Cat# 17101015 |
| Phalloidin-Alexa 647 | Thermo Fisher | Cat# A22287 |
| BODIPY 493/503 | MedChemExpress | Cat# HY-W090090 |
| Gentamycin | MedChemExpress | Cat# HY-A0276A |
| Vancomycin | MedChemExpress | Cat# HY-B0671 |
| BAPTA-AM | MedChemExpress | Cat# HY-100545 |
| Evans Blue | MedChemExpress | Cat# HY-B1102 |
| BHI broth | MERCK | Cat# 53286 |
| Puromycin | InvivoGen | Cat# ant-pr |
| Critical commercial assays |  |  |
| Peripheral blood mononuclear<br>Cell Isolation Kit | Solarbio | Cat# P8680 |
| Pierce Primary Neuron<br>Isolation Kit | Thermo Fisher | Cat# 88280 |
| Mouse IL-6 Uncoated ELISA<br>Kit | Thermo Fisher | Cat# 88-7064-77 |
| Mouse TNF-α Uncoated<br>ELISA Kit | Thermo Fisher | Cat# 88-7324-88 |
| CyQUANT LDH<br>Cytotoxicity Assay Kit | Thermo Fisher | Cat# C20301 |
| β-galactosidase Assay Kit | Thermo Fisher | Cat# 75707 |
| Glucose Colorimetric<br>Detection Kit | Thermo Fisher | Cat# EIAGLUC |
| Acid Phosphatase Assay | Abcam | Cat# ab83367 |
| β-hexosaminidase Assay Kit | Nanjing Jiancheng Bioengineering Institute | Cat# H485-1-2 |
| ATP Assay Kit | Sigma-Aldrich | Cat# MAK473 |
| ADP/ATP Ratio Assay Kit | Sigma-Aldrich | Cat# MAK135 |
| Lactate Assay Kit | Sigma-Aldrich | Cat# MAK570 |
| HSL ELISA Kit | Mlbio | Cat# ml026143 |
| Hematoxylin and Eosin Staining Kit | Beyotime | Cat# C0105S |
| Oil Red O Staining Kit | Beyotime | Cat# C0157 |
| Non-endotoxin plasmid small extraction medium dose kit | TIANGEN | Cat# DP118 |
| HiScript II 1st Strand cDNA Synthesis Kit | Vazyme | Cat# R212 |
| XF Glycolysis Stress Test kit | Seahorse Biosciences | Cat# 103020-100 |
| Experimental Models: Organisms/Strains |  |  |
| Mouse: C57BL/6J | Charles River | Cat# 219 |
| Recombinant DNA |  |  |
| pLV3-CMV-Chmp4b(mouse)-3×FLAG-CopGFP-Puro | Miaoling Biology | Cat# P44204 |
| Software and algorithms |  |  |
| GraphPad Prism 9 | GraphPad Software | https://www.graphpad.com |
| ImageJ | NIH | https://imagej.nih.gov/ij/ |
| FlowJo V10 | FlowJo LLC | https://www.flowjo.com |
| R studio | r-project.org |  |
| Other |  |  |
| Ketone ester | Guangao Biotechnology | Cat# GA8901 |
| Ketogenic diet (KD) | Xietong Pharma | Cat# XTKD01 |

### Clinical samples

Patients diagnosed with bacterial CNS infection (n = 62, 32 males and 30 females, aged 41.62 ± 15.68 years) and healthy donors (n = 60, 30 males and 30 females, aged 38.32 ± 9.17 years) were recruited from Chongqing Public Health Medical Center.

Patients were all exposed to *Mycobacterium tuberculosis* without immunosuppression. Patients were gender- and age-matched and were not concurrently infected with other pathogens, such as influenza or HIV. Meanwhile, subjects were recruited without any metabolic disorders, such as obesity or diabetes, in order to exclude their direct effects on BHB and CD36^+^ macrophages levels. Since blood glucose is a risk factor for bacterial neuroinvasion, subjects with abnormal blood glucose or insulin concentrations were also excluded. Clinical symptoms exhibited by the patients included a state of exhaustion, headaches, irritability, vomiting, fever, neck stiffness, convulsions, and consciousness disorder. Each symptom was assigned a value of one point. The clinical data of the patients were collected within 12 hours of admission.

The entire process of seven hospitalized patients from admission to recovery and discharge was followed up. Blood samples were collected and CD36^+^ macrophages and BHB levels were detected in all seven patients within 12 hours of admission, and changes in the CD36^+^ macrophages and BHB levels were measured continuously throughout the hospitalization period. According to the medical records, the entire hospitalization period was divided into a symptomatic period and a convalescent period, based on the recovery time points at which indicators of CNS infection (including white blood cell counts, protein levels and bacterial counts in the cerebrospinal fluid), and clinical symptoms of the patients returned to normal. All data were normalized to the parameters of first detection on admission.

Isolation of PBMCs and single-cell suspensions was performed by density gradient centrifugation using PBMC Isolation Kit. Plasma from each patient and donor was collected and then used for quantification of BHB. This study was reviewed and approved by the Ethics Committee at the Chongqing Public Health Medical Center, and was compliant with the “Guidance of the Ministry of Science and Technology (MOST) for the Review and Approval of Human Genetic Resources”. Healthy individuals and patients providing blood samples were given informed consent. The ethics committee approved this consent procedure.

The data in retrospective study were obtained from the Medical Information Mart for Intensive Care IV (MIMIC-IV) version 1.0 (https://physionet.org/content/mimiciv/1.0/), which includes medical information on patients in the intensive care unit of a hospital in Boston, USA, over a 10-year study period. The database was established with the approval of the Massachusetts Institute of Technology (Cambridge, MA) and the Institutional Review Boards of Beth Israel Deaconess Medical Center (Boston, MA). Data were extracted from the MIMIC-IV database using Structured Query Language (SQL) with Navicat Premium (version 12.0.28). A total of 128 admission records (49 patients /79 controls) from the MIMIC IV database were included in this study. Patients with benign intracranial hypertension were considered as controls.

### Animals

Animal experimental procedures were approved by the Laboratory Animal Welfare and Ethics Committee of Chongqing University. All mice were housed in a specific pathogen-free (SPF) facility according to standard humane animal husbandry protocols. In all experiments, experimental and control mice were 6-8-week-old littermates of C57BL/6 mice. HCAR2^-/-^, PCSK9^-/-^, HMGCS2^-/-^ mice and littermate controls obtained from heterozygote crosses were used for all experiments. Each in vivo experiment was performed using 50% female and 50% male animals, and results were not expected to be influenced by sex. All mice were maintained under a 12-hour light-dark cycle with ad libitum access to regular food and water.

### Bacterial strains

All procedures related to pathogenic bacteria were conducted in accordance with Biosafety Level 2 protocols and guidelines. The *L. monocytogenes* (ATCC 19115) and *S. pneumoniae* (ATCC 49619) were grown in Brain Heart Infusion Broth. The Methicillin-resistant *Staphylococcus aureus* USA300 (ATCC TCH1516) and *S. Typhimurium* (ATCC SL1438) were grown in Luria-Bertani Broth. Bacteria were grown at 37℃ with 5% CO_2_ to mid-log phase, and growth was evaluated by monitoring at an optical density of 600 nm. Frozen stocks of bacteria in 20% glycerol were prepared and kept at −80℃ until use.

### Data and code availability

Single-cell RNA-seq data (accession number GEO: GSE264020, secure token for reviewer: stchicgorhozbkb) and RNA-seq data (accession number GEO: GSE273820, secure token for reviewer: gbohyqeanrmzhyn; accession number GEO: GSE283713, secure token for reviewer: gbujwcoejdchtwz) have been deposited in GEO and are publicly available as of the date of publication.

### Mouse model of acute neuroinvasion

Mice were injected intravenously of 1×10^7^ CFU or intraperitoneally of 2×10^8^ CFU *L. monocytogenes* to establish an acute neuroinvasion mouse model. Gentamicin (30 mg/kg) was administered intraperitoneally 4 hours post infection to eliminate circulating bacteria that did not enter the phagocytes. Mice challenged with LPS (2 mg/mouse) served as positive controls for BBB damage. For other intracellular bacteria induced neuroinvasion models, mice were infected with MRSA (2×10^7^ CFU intravenously or 2×10^8^ CFU intraperitoneally), *S. Typhimurium* (1×10^7^ CFU intravenously or 1×10^8^ CFU intraperitoneally), or *S. pneumonia* (1×10^7^ CFU intravenously or 1×10^8^ CFU intraperitoneally), respectively. In the adoptive macrophages transfer experiments, bone marrow irradiation-based immunosuppression resulted in mice being more susceptible to bacterial infection, which necessitated a reduction of the infectious dose. After recovery from transfer for 12 hours, recipient mice were challenged with *L. monocytogenes* (1×10^6^ CFU intravenously), MRSA (2×10^6^ CFU intravenously), or *S. pneumonia* (1×10^6^ CFU intravenously) unless otherwise indicated.

The progression of acute bacterial meningitis was determined by clinical scoring as previously described with adjustments.^65^ Clinical scoring was performed based on the Table S2, including activity (0-2), time to return to upright position (0-2), eyes (0-2) and neurological examination (0-2). Mice were deeply anaesthetized by intraperitoneal injection of tribromoethanol solution (Avertin, 500 mg/kg), and perfused transcranially with saline solution for brain tissue pathomorphology. Blood was sampled immediately before perfusion. Tissue samples, including brain, lung, liver, and spleen, were dissected and weighed.

### Adoptive macrophage transfer experiments

Primary peritoneal macrophages from donor mice were collected 72 hours after intraperitoneal injection of thioglycolate. To collect CD36^+^ macrophages, purified macrophages were stimulated with 80 μg/ml ox-LDL for 24 hours and then sorted by an APC-conjugated CD36 antibody. Recipient mice were weighed 25-28 g and anesthetized for irradiation (Co 60, 10 Gy). 12 hours later, adoptive macrophages transfer was performed. After recovery for another 12 hours, recipient mice were challenged with bacteria (see mouse model of acute neuroinvasion section for infectious routes and doses), and extracellular bacteria were eliminated by gentamicin (30 mg/kg) or vancomycin (10 mg/kg) treatment 4 hours post infection. Samples were collected at 24 hours post infection for subsequent examinations unless otherwise indicated.

### Mouse diet Study and glucose supplementation

Mice were administered a supplement of 20 mg/ml ketone ester (D-β-hydroxybutyrate-(R)-1,3 butanediol monoester) in the drinking water for 4 weeks before the infection. The ketogenic diet contained 10% protein and 90% fat, whereas the control diet contained 23.2% protein, 2.8% fat and 74% sugar. Mice were placed on either the ketogenic or control diet for an additional 4 weeks before infection.

The food consumption was estimated by a subtraction method with the automated weighing scales as previously reported.^66^ A computer analyzed the changes in weight of each cage and outputs data on food intake by blocks of hours. The value of consumption per mouse was the amount of food reduced per cage divided by the number of mice per cage.

In the glucose supplementation experiment, mice were injected intraperitoneally with saline or glucose (100 mg/kg) every 12 hours after infection. To measure blood glucose levels, a drop of blood was collected by snipping the very end of the tail, and the blood in the strip was analyzed using a glucometer to measure blood glucose levels.

### Analysis of bacterial load in cells or tissues

Tissue samples of mice were weighted and then transferred into 2 ml tubes containing 4 mm stainless steel beads and 1 ml of ice-cold sterile saline. Homogenization was carried out in TissueLyser II (Qiagen) for 5 minutes at 70 Hz. Serial dilutions of the homogenates were plated on the plates incubated at 37℃ and CFUs were counted. The infected macrophages were incubated in fresh solution containing 5 μg/mL lysostaphin for the indicated periods and then lysed by 0.2% Triton to release intracellular bacteria. The lysates were diluted to appropriate dilutions with sterile PBS for CFU count on agar plates.

### BBB permeability assays

After infection, mice were retro-orbitally injected with 2% Evans blue. After 2 hours of circulation, mice were deeply anaesthetized, and the brains were delicately dissected after transcardial PBS perfusion. Whole-brain images were acquired using a digital camera. To quantify Evans blue leakage, mouse brains were sectioned longitudinally, weighted and then homogenized in 0.5 ml of 50% trichloroacetic acid. Lysates were centrifuged at 15,000 g for 30 minutes. The amount of extravasated Evans blue in the supernatant was measured at 620/680 nm emission using a Multi-Mode Microplate Reader (SpetraMax i3x), calculated by a standard curve, and normalized to the weight of each brain.

### Brain histopathology and Immunostaining

Brain samples were collected from infected mice 24 hours after injection. Mice were deeply anaesthetized with tribromoethanol solution (Avertin, 500 mg/kg, intravenously), followed by transcranial perfusion with 30 ml of saline solution, and then of 4% Paraformaldehyde (PFA) in 10 ml of saline. After dissection, brain samples were maintained in 4% PFA at 4℃. Fixed brains were embedded in paraffin, sectioned and half of the slides were stained using haematoxylin and eosin. Each field was assigned a score from 0 to 3 based on neuronal morphology: grade 0 (no alteration); grade 1 (no vacuolation with small numbers of pyknotic cells); grade 2 (moderate vacuolation and pyknosis); and grade 3 (extensive vacuolation, pyknosis and tissue loss or liquefactive necrosis).

Fresh frozen brain tissue sections of 5-10 mm thickness were mounted on slides, air dried for 10 minutes at room temperature, and fixed in ice cold 10% neutral-buffered formalin for 5-10 minutes. Slices were washed in PBS, incubated in 3% bovine serum albumin (BSA) for 2 hours, and then washed with 0.1% Triton X-100 in PBS. The slides were labeled with the appropriate primary antibodies overnight at 4℃ in mild blocking conditions. Then next day, the slices were washed in PBS, and then incubated with secondary antibodies and DAPI. Tissues were washed in PBS and then mounted on glass slides under coverslips in mounting medium. The slides were left in darkness overnight before observation under a Zeiss LSM900 or LM710 microscope.

### Cell sorting and flow cytometry

Isolated mouse PBMCs were washed twice with PBS. The proportion of CD36^+^ macrophages population was determined by adding appropriate amount of fluorescent-coupled primary antibodies (F4/80-PE, CD11b-FITC, CD36-APC). The cells were incubated at 4℃ for 30 minutes away from light. The living macrophages were collected according to the expression of CD36 and then cultured overnight *in vitro*. The lipid content in sorting cells was detected using BODIPY FL and oil Red O. To evaluate the lipid uptake capacity, macrophages were incubated with 40 µg/ml Dil-ox-LDL particles for 6 hours, and then the fluorescence intensity was measured by flow cytometry. To detect the expression of LAMP1 on the cell surface, infected macrophages were washed twice with PBS and then stained with LAMP1-FITC at 4℃ for 30 minutes away from light. The cells were washed three times and then re-suspended in 200μl of the staining buffer for final FACS analysis. Data was analyzed with FlowJo software.

### Isolation and culture of primary cells

To generate mouse peritoneal macrophages, 8-week-old mice were injected intraperitoneally with 3% thioglycolate broth. After 72 hours, peritoneal cells were harvested and macrophages were enriched by centrifuge. To obtain primary mouse brain microvascular endothelial cells (BMECs), the cerebral cortex in the brains were separated and rolled on the filter paper to remove the meninges and choroid plexuses. A mechanical reduction was performed by mincing the brain pieces using surgical blades. The homogenate was centrifuged (800 g, 8 minutes, 4℃) and the precipitate was digested in DMEM containing 1 mg/ml collagenase II and 58.5 U/ml DNase I for 75 minutes on a benchtop orbital shaker at 37℃. The supernatant was collected and centrifuged at 500 g for 8 minutes. The isolated cerebrovascular endothelial cells were purified by the anchoring technique. The livers and lungs from anaesthetized mice were perfused via the portal vein sequentially with balanced salt solution containing 5 mM glucose, 0.01% sodium heparin, and 5 mM EGTA and DMEM medium containing 4 mM CaCl_2_, 2% FBS and 0.05% collagenase IV at a flow rate of 5 ml/min for 5 minutes, respectively. The following steps are similar in order to separate the liver sinusoids endothelial cells and pulmonary vascular endothelial cells. Isolation of primary embryonic neurons and single-cell suspensions was performed using Pierce Primary Neuron Isolation Kit by density gradient centrifugation.

### *In vitro* blood-brain barrier model

The *in vitro* BBB model was constructed with primary endothelial cells using a Transwell cell culture system. Briefly, BMECs were seeded onto the upper chamber of a Transwell pre-coated with gelatine, and cultured with the medium containing 10% FBS. The integrity of the monolayer cell was evaluated by measuring the TEER values using a Millicell-ERS voltohmmeter (Millipore). The cell monolayers with TEER values higher than 300 Ω.cm^2^ were used as BBB models for the transmigration studies. The neurons cells were seeded onto the lower chamber of the Transwell for bacterial loading.

Macrophages were exposed to *L. monocytogenes* (MOI=10) for 6 hours and then treated with 100 µg/ml gentamicin for 2 hours to kill the extracellular bacteria. After construction of the BBB model, the infected macrophages were trypsinized for cell counting and added into the upper chamber. The incubation time was 12 hours (unless otherwise indicated) and then the neurons cells in the bottom chamber were lysed for penetrating bacterial quantification.

### Live-cell flow chamber assays

The adhesion, shear resistance, crawling, and transmigration of macrophages on endothelial monolayer were tested using a flow cell system as previously described.^40^ Shear flow was applied by a syringe pump (PHD22/2000, Harvard Apparatus) into a microfluidic chip with a 100 μm × 1 mm × 15 mm main flowing zone. Different shear stresses were acquired by setting corresponding volumetric flow rates. Isolated macrophages were cultured with different concentrations of ox-LDL for 24 hours and then infected with *L. monocytogenes* (MOI=10) for 6 hours. Macrophages were re-suspended and stained with Hoechst. The cells were then perfused over pre-formed endothelial monolayer at a shear stress of 0.1 dyn/cm^2^ for 8 minutes. Then, the adhesion and crawling dynamics of macrophages on endothelial monolayer were recorded at shear stress of 2 dyn/cm^2^ for 10 minutes, by using a CCD camera with 20× objective. Images were taken from the video every second and analyzed for the crawling parameters and transmigrating duration with Image J software.

The post-arrest crawling of macrophages on the microfluidic chip can be factorized into two metrics: the forward migration index (FMI) towards the x-axis (along with flow, xFMI), and the y-axis (perpendicular to flow, yFMI). xFMI and yFMI were accurately measured by a CCD camera, and these two indices constituted the crawling trajectories of each individual cell shown in Figure 3E. Directionality was quantified as the ratio of yFMI to xFMI. Directionality indicates the angle at which the direction of cell movement deviates from the shear flow. The larger the value of directionality, the less the direction of cell movement is affected by the shear flow, indicating that the cell has a stronger ability to fight against the shear flow. Transmigration duration, which is defined as the interval from the timepoint of cell arrival at the site to the endpoint of completion of transmigration beneath the endothelial monolayer (Figure S6H), was used to indicate the transmigration capacity of the cell. Here a macrophage crossing the monolayer with a 70% fluorescence intensity reduction was deemed transmigrated.

For testing shear-resistant macrophage adhesion, primed macrophages were re-suspended and counted. The same number of cells (2×10^4^) was completely perfused over pre-formed endothelial monolayer within 10 minutes. The shear stress was increased stepwise from 1 to 2, 4, 8, 16, 32 dyn/cm^2^ for 30 seconds each. Remaining adherent macrophages on endothelial monolayer were recorded at 20× objective by a CCD camera and counted using Image J software. The normalized adhesion fraction was calculated by dividing the number of adherent macrophages at the endpoint of each shear phase by the number at the endpoint of 1 dyn/cm^2^. For defining the macrophage transmigration dynamics, the time-lapse images were analyzed offline frame-by-frame every second.

### Quantification of cell area and protrusions

The cell area and protrusions of macrophages were measured using a quantitative cell morphology analysis system adapted from Young et al.^67^ In brief, peritoneal macrophages were stimulated with different concentrations of ox-LDL for 24 hours, and then cells were fixed, permeabilized, and labeled with phalloidin-Alexa 647. Samples were analyzed using a Zeiss LSM900 confocal microscope. 8-bit images were acquired by using ImageJ software and converted to grayscale to best visualize all positive staining. Brightness and contrast were adjusted and a despeckle step was performed as needed until cell edges were clearly visible. For fractal analysis, images were skeletonized using the plugin AnalyzeSkeleton (2D/3D). Skeleton fragments generated during image acquisition were removed to facilitate quantification. Protrusions were identified as the actin tails extending from the cells surface. The area of the cells as well as the length of the actin tails were measured using ImageJ software. At least ten cells were scored per group.

### Single cell RNA-sequencing analysis

PBMCs isolated from mice were filtered using a 40-μm nylon cell strainer and red blood cells were removed by red blood cell lysis solution. Dissociated cells were washed with DPBS containing 2% FBS. Cells were stained with 0.4% Trypan blue to check the viability. Sequencing libraries were prepared using randomly interrupted whole-transcriptome amplification products to enrich the 3’ end of the transcripts linked with the cell barcode and unique molecular identifier. All the procedures including the library construction were performed strictly according to the standard manufacturer’s protocol [10× Genomics Chromium Single-Cell 3′ kit (V3)]. Sequencing libraries were quantified using a High Sensitivity DNA Chip on a Bioanalyzer 2100 and the Qubit High Sensitivity DNA Assay. Single cells were sequenced using an Illumina NovaSeq 6000 sequencing system (paired-end multiplexing run, 150 bp) by Majorbio Bio-pharm Technology Co., Ltd (Shanghai, China). The sequencing and bioinformatic analysis were performed on platform of Majorbio (Shanghai, China).

Reads were processed using the Cell Ranger (v7.1.0) pipeline with default and recommended parameters. FASTQs generated from Illumina sequencing output were aligned to the mouse genome, version GRCm38, using the STAR algorithm. Gene-barcode matrices containing the barcoded cells and gene expression counts were generated, and then imported into the Seurat (v3.2.0) R toolkit for quality control and downstream analysis of our single cell RNAseq data.

Cells were clustered using graph-based clustering of the PCA reduced data with the Louvain Method after computing a shared nearest neighbor graph. For sub-clustering, we applied the same procedure of scaled, dimensionality reduction, and clustering to the specific set of data. For each cluster, we used the Wilcoxon Rank-Sum Test to find genes with significant deferential expression compared to the remaining clusters.

Differentially expression genes (DEGs) were identified with Seurat’s FindMarkers, using a likelihood ratio test. DEGs with |log_2_ (fold change)| > 1 and p value ≤ 0.05 were considered to be significant. In addition, GO functional-enrichment analysis was performed to identify which DEGs were significantly enriched in GO terms and metabolic pathways at Bonferroni-corrected P-value ≤ 0.05 compared with the whole-transcriptome background.

### Lipid metabolomics

Metabolite extraction and mass spectrometry was performed using an Ultra-High Performance Liquid Chromatography-mass spectrometry (UHPLC-MS) by Biomarker Technologies (Beijing, China). The raw data was collected under the control of the acquisition software (MassLynx V4.2, Waters), and then processed by Progenesis QI software integrated with the METLIN database (Waters). The theoretical fragment identification and mass deviation were within 100 ppm. Following normalization of peak areas relative to the total peak area, principal component analysis (PCA) and Spearman correlation analysis were performed to evaluate the repeatability of the samples. A total of 4,625 named metabolites were identified for the datasets. The identified compounds were classified, and pathway information was annotated by using the KEGG, HMDB and LipidMaps databases. The enrichment significance of differential metabolites in KEGG pathways was evaluated using a hypergeometric distribution test.

### RNA-seq (RNA Sequencing)

RNA was extracted from mouse liver tissues using a Trizol extraction and Purelink RNA kit (ThermoFisher Scientific). Bioanalyzer chips were used to assess RNA quality prior to RNA sequencing. A total of 1 μg RNA per sample was used as input material for RNA sample preparation. Sequencing libraries were generated using a Hieff NGS Ultima Dual-mode mRNA Library Prep Kit for lumina following manufacturer’s recommendations, and index codes were added to attribute sequences for each sample. The libraries were sequenced on an Illumina NovaSeq platform to generate 150 bp paired-end reads, according to the manufacturer’s instructions.

### Measurement of ECAR

ECAR was measured using the XF Glycolysis Stress Test kit. In brief, liver tissue was inoculated into XFe24 Cell Culture Microplates and cultured overnight at 37℃ in an incubator containing 5% CO_2_. Three injections (10 mM glucose, 1.5 μM oligomycin and 50 mM 2-DG) were performed at the indicated timepoints. ECAR values were normalized to the total amount of proteins. ECAR parameters were calculated using Wave software (v.2.4.3, Seahorse Bioscience, Agilent Technologies) according to manufacturer instructions.

### Fura-4 Ca^2+^ imaging

Cells were washed with PBS and incubated with 5 μM Fluo-4 AM and 0.04% Pluronic-127 for 60 minutes at 37℃. Then, cells were washed with PBS and incubated with PBS for 30 minutes at 37℃. Fluorescence was detected using a CytoFLEX Flow Cytometer (Beckman Coulter) equipped with a 488 nm excitation wavelength. 30 seconds after loaded with Fluo-4 AM, cells were then treated with BHB (10 mM) as indicated in the figures. Data was analyzed with Flow Jo to determine the fluorescence intensity.

### Real-time qRT-PCR

Total RNA was isolated using a RNeasy Mini Kit (QIAGEN). The mRNA was reverse-transcribed and quantified using a HiScript II 1st Strand cDNA Synthesis Kit (Vazyme). qRT-PCR was performed on an ABI StepOnePlus PCR system (Thermo Fisher), and results were normalized to GAPDH mRNA levels. Data were analyzed using the 2^-ΔΔCT^ method. All the primers used for qRT-PCR are listed in Table S3.

### Statistical analysis

Statistical analysis was performed using GraphPad Prism Software. Two-group comparisons were made using two-tailed unpaired Student’s t-test. One-way analysis of variance (ANOVA) with appropriate multiple comparisons tests was used to compare three independent groups. For comparisons of multiple factors, two-way ANOVA with appropriate multiple comparisons tests was used. A value of p < 0.05 was considered significant. Specific details of statistical analysis are presented above or in associated figure legends.

---

**Figure S1.**
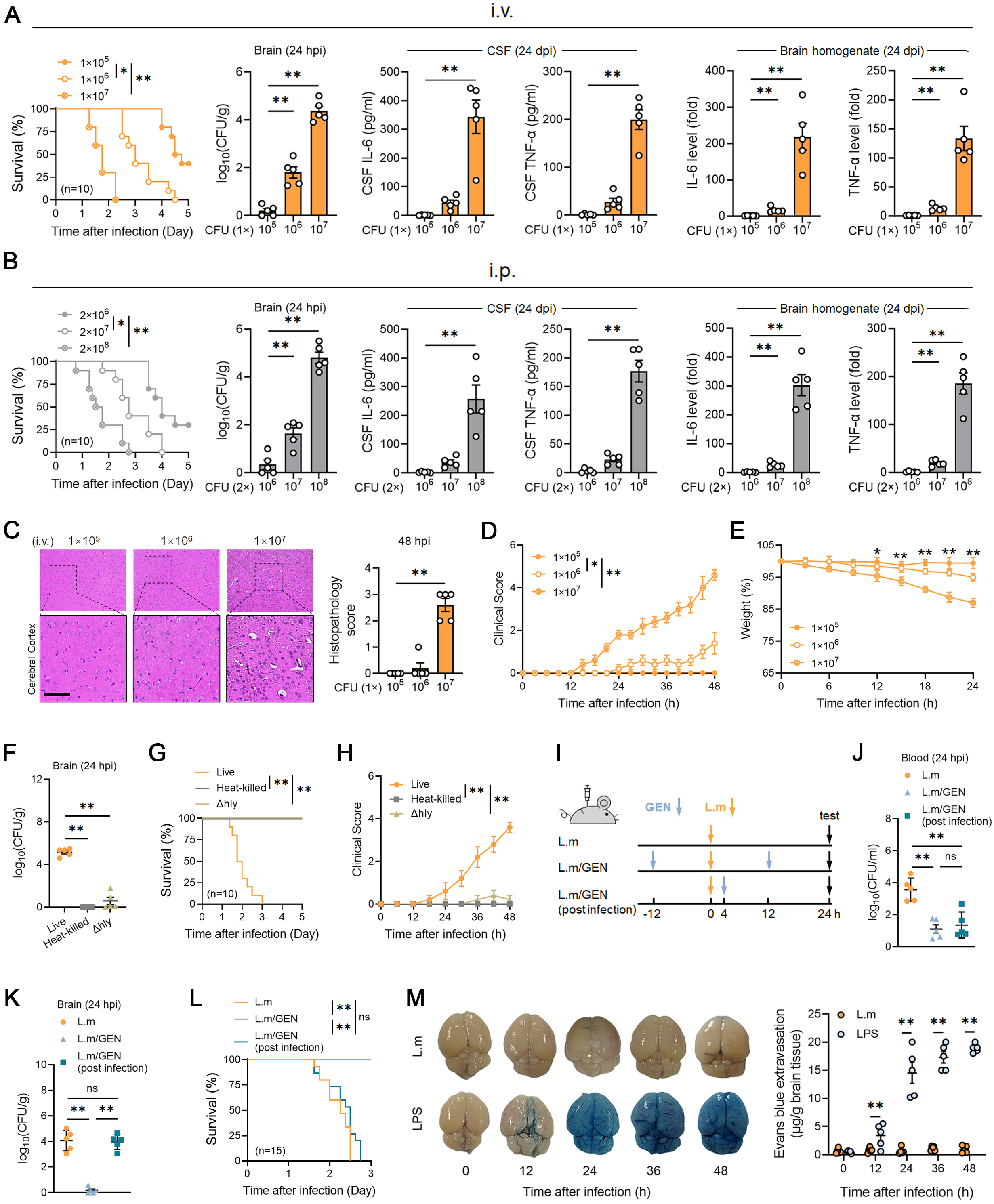
Acute bacterial neuroinvasion mouse model induced by *L. monocytogenes*. (A) Mice were intravenously infected with 1×10^5^, 1×10^6^, or 1×10^7^ CFU of *L. monocytogenes*. Survival, bacterial burdens, and pro-inflammatory factor levels in the CSF and brain homogenates were evaluated. (n=10 for each group) (B) Mice were intraperitoneally infected with 2×10^6^, 2×10^7^, or 2×10^8^ CFU of *L. monocytogenes*. Survival, bacterial burdens, and pro-inflammatory factor levels in the CSF and brain homogenates were evaluated. (n=10 for each group) (C-E) Mice were intravenously infected with 1×10^5^, 1×10^6^, or 1×10^7^ CFU of *L. monocytogenes*. (C) Representative images of HE staining of the cerebral cortex tissues and pathology scores. (D) Clinical scores. (E) Body weight changes. (F-H) Mice were intravenously infected with 1×10^7^ CFU of live, heat-killed or attenuated (Δhly) *L. monocytogenes*. (n=10 for each group) (F) Bacterial burdens of brains. (G) Survival rate. (H) Clinical scores. (I-J) (I) Experimental diagram of mice treated with gentamicin. Mice were intravenously infected with 1×10^7^ CFU of *L. monocytogenes*. Gentamicin (30 mg/kg) was administered to mice daily during the infection or 4 hours post-infection. (n=15 for each group) (J-K) Bacterial burdens in the peripheral blood (J) and brain tissues (K). (L) Survival of the mice. (M) Representative images and quantification of Evans blue leakage in mouse brains. Evans blue assay was performed on mice challenged with *L. monocytogene*s (1×10^7^ CFU intravenously) or LPS (2 mg/mouse). Experiments were performed in duplicate. Data are presented as mean ± SEM (n=5). *p < 0.05, **p < 0.01. Abbreviations: L. m, *L. monocytogene*; i.v., intravenous; i.p., intraperitoneal; hpi, hours post infection; GEN, gentamicin.

**Figure S2.**
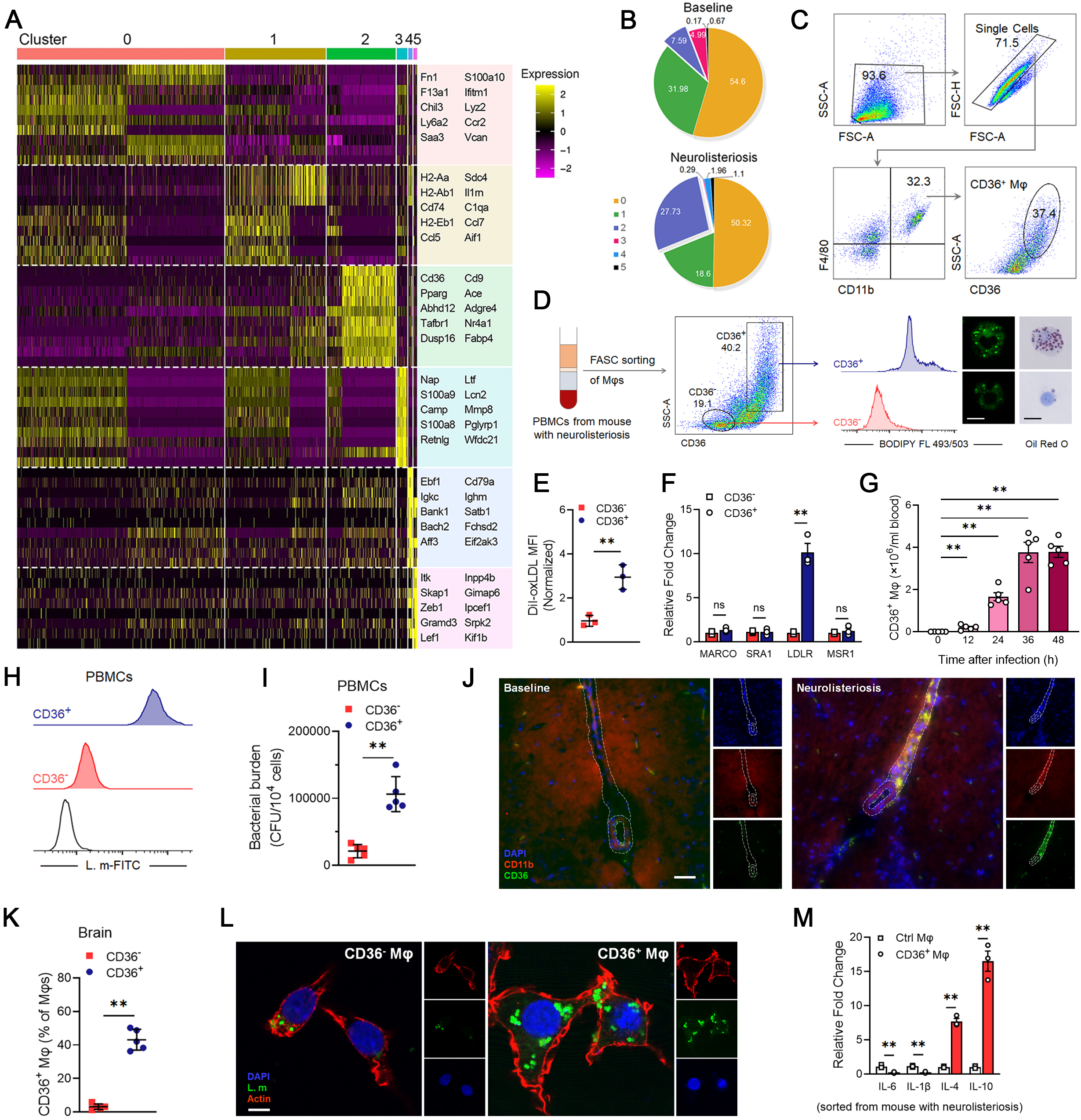
Additional results of identification of CD36^+^ macrophages during neurolisteriosis, related to Figure 1. (A) Cell clusters identified in 10× scRNA-seq dataset. The top 10 enriched genes for each cluster are highlighted. Normalized z-score is used to illustrate gene expressions. (B) Proportion of each cluster according to experimental conditions in Figure 1D. (C) Gating strategy for analysis of CD36^+^ macrophages in PBMCs. (D-F) (D) FACS of the CD36^+^ and CD36^-^ macrophages from peripheral blood and staining of BODIPY FL 493/503 and oil Red O. Scale bar, 10 μm. Quantification of the normalized intake of Dil-oxLDL (E) and mRNA expressions of the scavenger receptors (F) in sorted CD36^+^ and CD36^-^ macrophages. (G) Quantification of the absolute numbers of CD36^+^ macrophages in PBMCs according to experimental conditions in Figure 1F. (H-I) Quantification of the intracellular bacterial burdens of the CD36^+^ and CD36^-^ macrophages sorted from peripheral blood by flow cytometry (H) and plate dilution method (I). (J) Immunofluorescence staining of CD36^+^ macrophages in cerebral vasculature of mice with neurolisteriosis. CD36^+^ macrophages were stained with CD11b (red) and CD36 (green). The boundary of vascular was depicted by dashed lines. Scale bar, 100 μm. (K-L) CD36^-^ and CD36^+^ macrophages were sorted from brain tissues of infected mice according to experimental conditions in Figure 1J. (K) Percentages of CD36^-^ and CD36^+^ macrophages in the brain. (L) Immunofluorescence staining of infected CD36^-^and CD36^+^ macrophages. *L. monocytogenes* (green) and actin (red) were stained. Scale bar, 10 μm. (M) MRNA levels of the pro-inflammatory factors in control macrophages and sorted CD36^+^ macrophages. Experiments were performed in duplicate (A-D, G-L) or triplicate (E-F, M). Data are presented as mean ± SEM [n=3 in (E-F, M), n=5 in (G, I-K)]. *p < 0.05, **p < 0.01. Abbreviations: PMφ, peritoneal macrophages; CD36^-^ Mφ, CD36^-^ macrophages; CD36^+^ Mφ, CD36^+^ macrophages.

**Figure S3.**
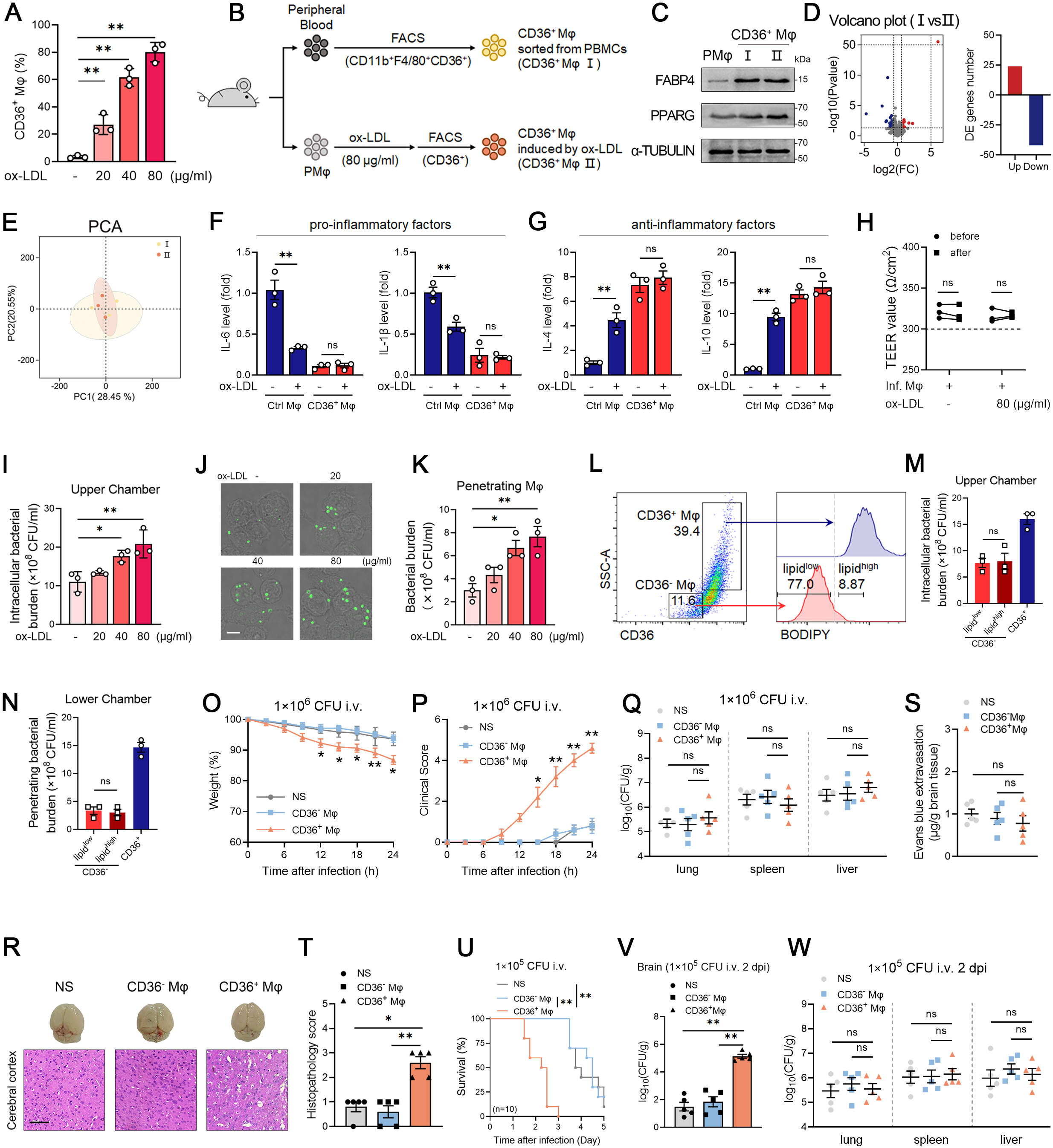
Additional results of adoptive transfer of CD36^+^ macrophages, related to Figure 2. (A) Mouse peritoneal macrophages were treated with different concentrations of ox-LDL for 24 hours, and then the percentage of CD36^+^ macrophages was assessed by flow cytometry. (B-E) (B) Schematic diagram of isolation of CD36^+^ macrophages from two different sources. (C) Immunoblot analysis of FABP4 and PPARG expressions. (D-E) Transcriptomics analysis of CD36^+^ macrophages from different sources. (n=3 for each group) (D) Differentially expressed genes (DEGs) were shown in the volcano plot. (E) PCA analysis of differences between groups. (F-G) MRNA expression of pro-inflammatory (F) and anti-inflammatory (G) factors in CD36^+^ and control macrophages sorted from PBMCs. (H-K) Experiments based on the *in vitro* BBB model shown in Figure 2A. (H) TEER values of the BBB model before and after bacterial infection. (I) Quantification of the intracellular bacterial burdens in macrophages in the upper chamber of Transwell system. (J-K) Representative images (J) and quantification (K) of the bacteria inside the macrophages that penetrating into the lower chamber. Scale bar, 10 μm. (L-N) (L) Gating strategy for sorting CD36^-^ lipid^high^ and CD36^-^ lipid^low^ macrophages in PBMCs. (M-N) Quantification of intracellular bacterial burdens in the upper chamber (M) and penetrating bacterial burdens in the neuron cells in the lower chamber of Transwell system (N). (O-W) Additional results of adoptive macrophage transfer experiments shown in Figure 2E. (O-T) Body weight changes (O). Clinical scores (P). Bacterial burdens in other organs of recipient mice at 24 hours post infection (Q). Representative images (R) of Evans blue leakage in mouse brains and HE staining in cerebral cortex tissues. Quantification of extravasation (S) and pathology scores (T). Scale bar, 20 μm. (U-W) After adoptive macrophage transfer, recipient mice were intravenously infected with 1×10^5^ CFU of *L. monocytogenes*. (n=10 for each group) Survival of recipient mice (U). Bacterial burdens in brains (V) and other organs (W) of recipient mice at 2 days post infection. Experiments were performed in duplicate (O-W) or triplicate (A, F-N). Data are presented as mean ± SEM [n=3 in (A, F-I, K, M-N), n=5 in (Q, S-T, V-W)]. *p < 0.05, **p < 0.01. Abbreviations: ns, nonsignificant; NS, normal saline; PMφ, purified peritoneal macrophages; CD36^-^ Mφ, CD36^-^ macrophages; CD36^+^ Mφ, CD36^+^ macrophages; ox-LDL, oxidized low densitylipoprotein; i.v., intravenous; i.p., intraperitoneal.

**Figure S4.**
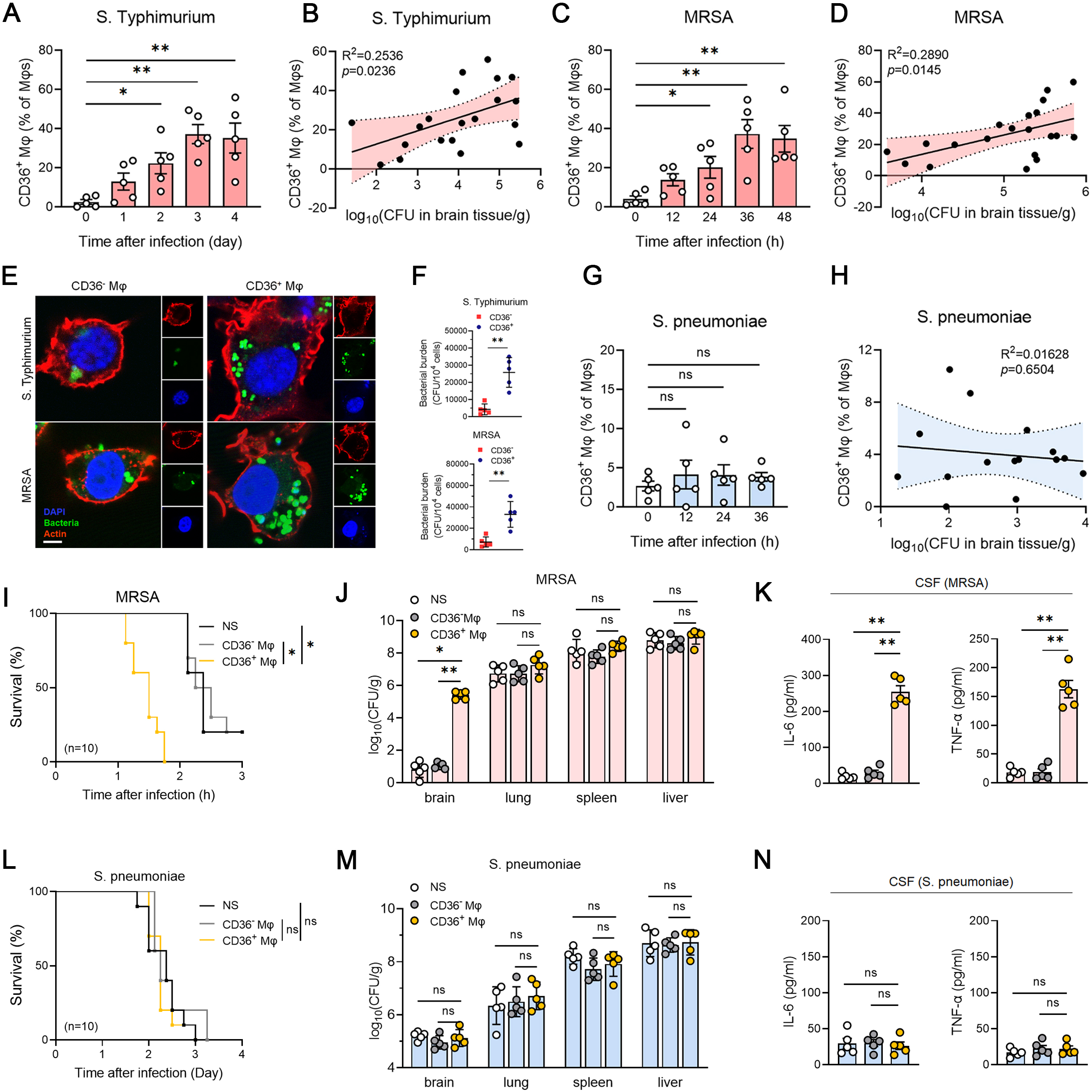
Accumulation of CD36^+^ macrophages exacerbates intracellular bacterial neuroinvasion. (A-H) Mice were intraperitoneally infected with 1×10^8^ CFU of *S. Typhimurium* (A-B), 2×10^8^ CFU of MRSA (C-D) or 1×10^8^ CFU of *S. pneumoniae* (G-H), respectively. (n=5 for each group) (A, C, G) Flow cytometry analysis of CD36^+^ macrophages during neuroinvasion. (B, D, H) Correlation between CD36^+^ macrophages and bacterial burdens in the brain during bacterial neuroinvasion. R, correlation coefficient. (E-F) CD36^-^ and CD36^+^ macrophages were sorted from brain tissues of mice infected with *S. Typhimurium* or MRSA. (E) Immunofluorescence staining of CD36^-^ and CD36^+^ macrophages. Bacteria (green) and actin (red) in macrophages were stained. Scale bar, 10 μm. (F) Quantification of the intracellular bacterial burden of the CD36^+^ and CD36^-^ macrophages by plate dilution method. (I-K) After adoptive macrophage transfer, recipient mice were intravenously infected with 2×10^6^ CFU of MRSA. Vancomycin (10 mg/kg) was administered intraperitoneally 4 hours post infection. (n=10 for each group) (I) Survival of recipient mice. (J) Bacterial burdens in tissues. (K) Pro-inflammatory factor levels in CSF of recipient mice on day 1 post infection. (L-N) After adoptive macrophage transfer, recipient mice were intravenously infected with 1×10^6^ CFU of *S. pneumoniae*. (n=10 for each group) (L) Survival rates of mice. (M) Bacterial burdens in tissues. (N) Pro-inflammatory factor levels in CSF of recipient mice on day 2 post infection. Experiments were performed in duplicate. Data are presented as mean ± SEM (n=5). *p < 0.05, **p < 0.01. Abbreviations: ns, nonsignificant; CD36^-^ macrophages; CD36^+^Mφ, CD36^+^ macrophages.

**Figure S5.**
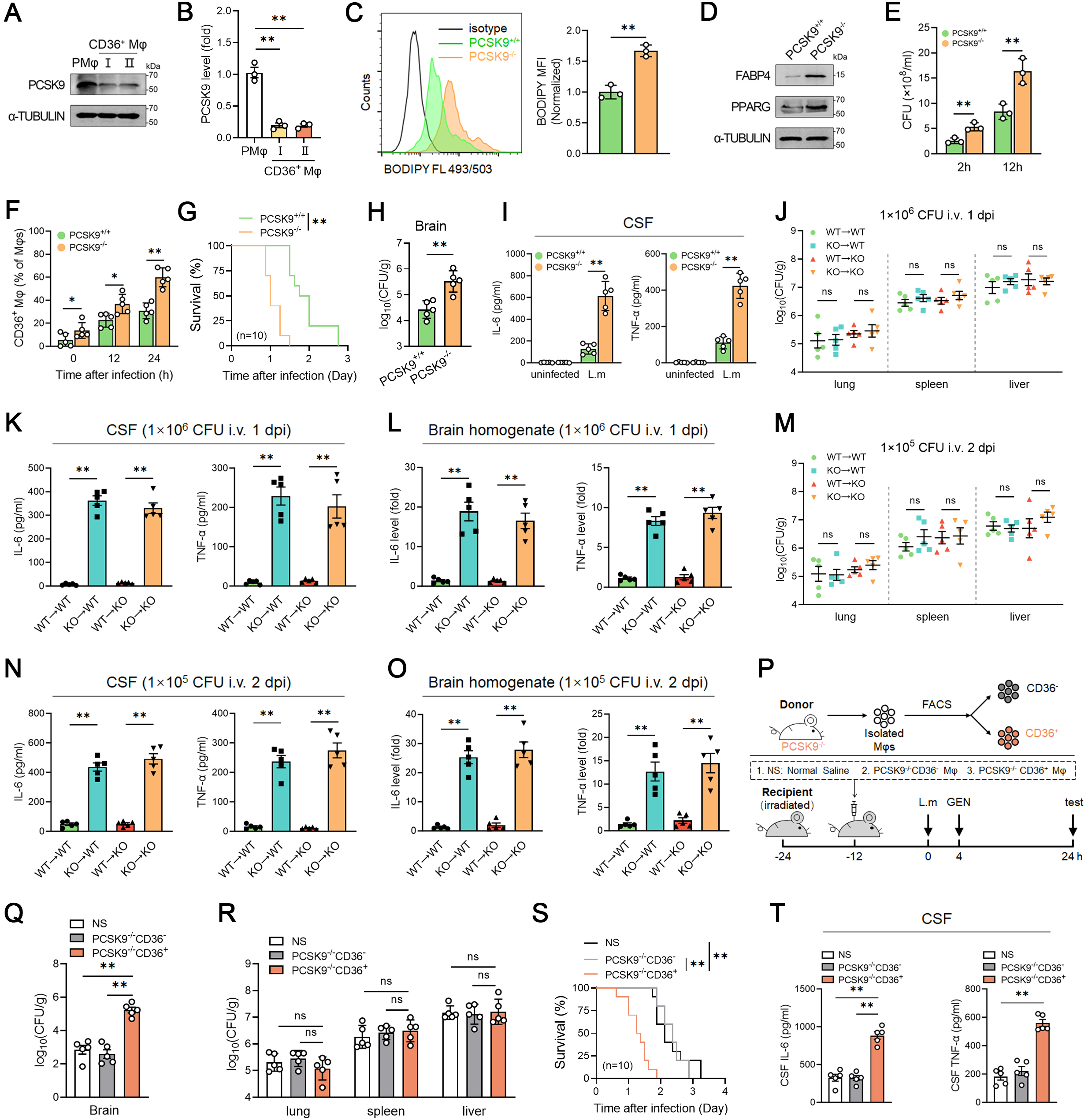
Additional results of bidirectional transfer of macrophages from PCSK9^+/+^ and PCSK9^-/-^ mice, related to Figure 2. (A-B) Immunoblot (A) and qRT-PCR (B) analysis of expression of PCSK9 in CD36^+^ macrophages. (C) Flow cytometry analysis of lipid droplets in peritoneal macrophages of PCSK9^+/+^ and PCSK9^-/-^ mice. (D) Immunoblot analysis of FABP4 and PPARG expressions. (E) Intracellular bacterial burdens of infected macrophages. Peritoneal macrophages of PCSK9^+/+^ and PCSK9^-/-^ mice were infected with *L. monocytogenes* (MOI=10) for 2 hours, and then cells were washed by PBS containing gentamicin. Intracellular bacterial burdens were calculated at 2 or 12 hours post infection. (F-I) PCSK9^+/+^ and PCSK9^-/-^ mice were intravenously infected with 1×10^7^ CFU of *L. monocytogenes*. (n=10 for each group) (F) Percentage of CD36^+^ macrophages by flow cytometry during neurolisteriosis. (G) Survival rates of mice. (H-I) Bacterial burdens in the brain (H) and pro-inflammatory factor levels in CSF (I) of mice at 24 hours post infection. (J-L) Additional results of adoptive macrophage transfer experiments shown in Figure 2K-2L. (J) Bacterial burdens in other organs (lung, spleen, and liver). (K-L) Pro-inflammatory factor levels in CSF (K) and brain homogenates (L) at day 1 post infection. (M-O) Additional results of adoptive macrophage transfer experiments shown in Figure 2M-2N. (M) Bacterial burdens in other tissues (lung, spleen, and liver). (N-O) Cytokine levels in CSF (N) and brain homogenates (O) on day 2 post infection. (P-T) (P) Schematic diagram of adoptive macrophage transfer experiments. PCSK9^-/-^ CD36^+^ and PCSK9^-/-^CD36^-^ macrophages were sorted from the peritoneal macrophages of PCSK9^-/-^ mice, and then injected into irradiated recipient mice. Gentamicin (30 mg/kg) was administered intraperitoneally 4 hours after infection. (n=10 for each group) (Q-R) Bacterial burdens in the brain (Q) and other organs (lung, spleen, and liver) (R). (S) Survival of recipient mice. (T) Pro-inflammatory factor levels in the CSF of recipient mice at 24 hours post infection. Experiments were performed in duplicate (F-T) or triplicate (A-E). Data are presented as mean ± SEM [n=3 in (B-C, E), n=5 in (F, H-O, Q-R, T)]. *p < 0.05, **p < 0.01. Abbreviations: ns, nonsignificant; i.p., intraperitoneally; dpi, days post infection; CD36^-^ macrophages; CD36^+^ Mφ, CD36^+^ macrophages.

**Figure S6.**
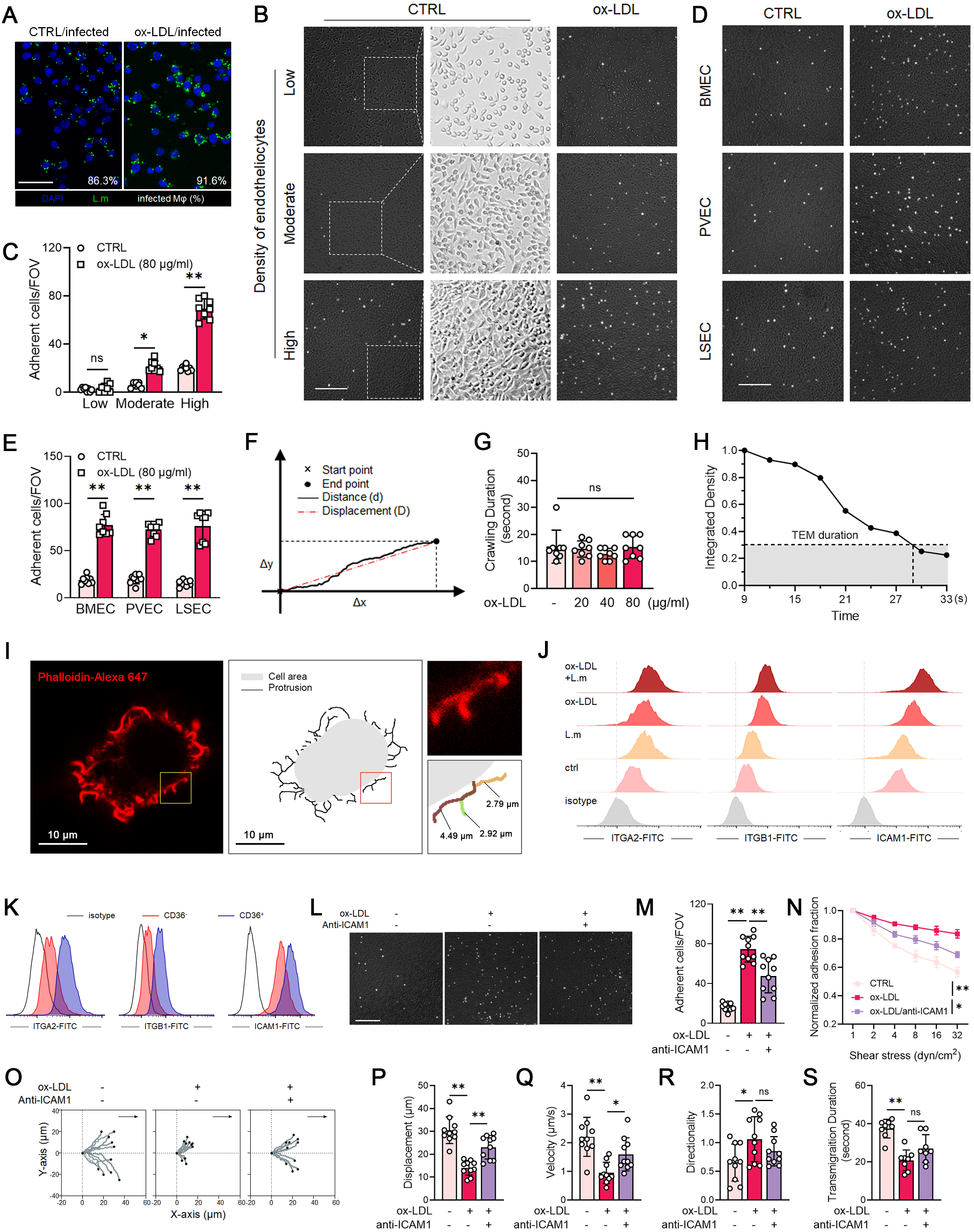
Supplemental data for the live-cell flow chamber assays, related to Figure 3. (A) Immunofluorescence staining of infected macrophages. Primary peritoneal macrophages were pretreated with ox-LDL for 24 hours and then infected with *L. monocytogenes* (MOI=10) for 6 hours. Intracellular bacteria (green) in macrophages were stained afterwards. Scale bar, 10 μm. (B-E) Optical images of *L. monocytogenes*-infected macrophages on the surface of endothelial monolayers with different densities (B) or endothelial monolayers derived from different tissues (D). Scale bar, 200 μm. (C, E) Quantification of adherent cells in each field of view. (F) Schematic trajectory used for defining macrophage crawling parameters. (G) Crawling duration of *L. monocytogenes*-infected macrophages after adhesion to endothelial monolayer surface. (H) Relative fluorescence intensity of a macrophage on endothelial surface over time. It is a random example to illustrate the definition of transmigration duration. A 70% decrease in fluorescence intensity after a macrophage crosses a monolayer is considered successful transmigration. (I) Representative images for quantifying the cell area and protrusions of macrophages. (J-K) Flow cytometry analysis of intercellular adhesion molecules in peritoneal macrophages (J) and sorted CD36^-^ and CD36^+^ macrophages (K). (L-S) Primary peritoneal macrophages were pretreated with ox-LDL for 24 hours and then infected with *L. monocytogenes* (MOI=10) for 6 hours. ICAM1 antibody (50 μg/mL) blockade was performed for 30 minutes prior to ox-LDL stimulation. (L) Optical images of adherent macrophages on the surface of monolayer endothelial cells. Scale bar, 200 μm. (M) Quantification of adherent cells in each field of view. (N) Macrophages resistance to shear flow after adhesion to endothelial cells. (O-S) Crawling and migrating dynamics of infected macrophages under shear flow. (O) Typical trajectories of infected macrophages crawling on monolayer endothelial cells. Quantification of crawling displacement (P), velocity (Q), directionality (R) and transmigration duration (S). Experiments were performed in triplicate. Data are presented as mean ± SEM [n=8 in (C, E, G), n=5 in (M, P-S)]. *p < 0.05, **p < 0.01. Abbreviations: BMEC, brain microvascular endothelial cell; PVEC, pulmonary vascular endothelial cell; LSEC, liver sinusoidal endothelial cell; TEM, transendothelial migration; ns, nonsignificant.

**Figure S7.**
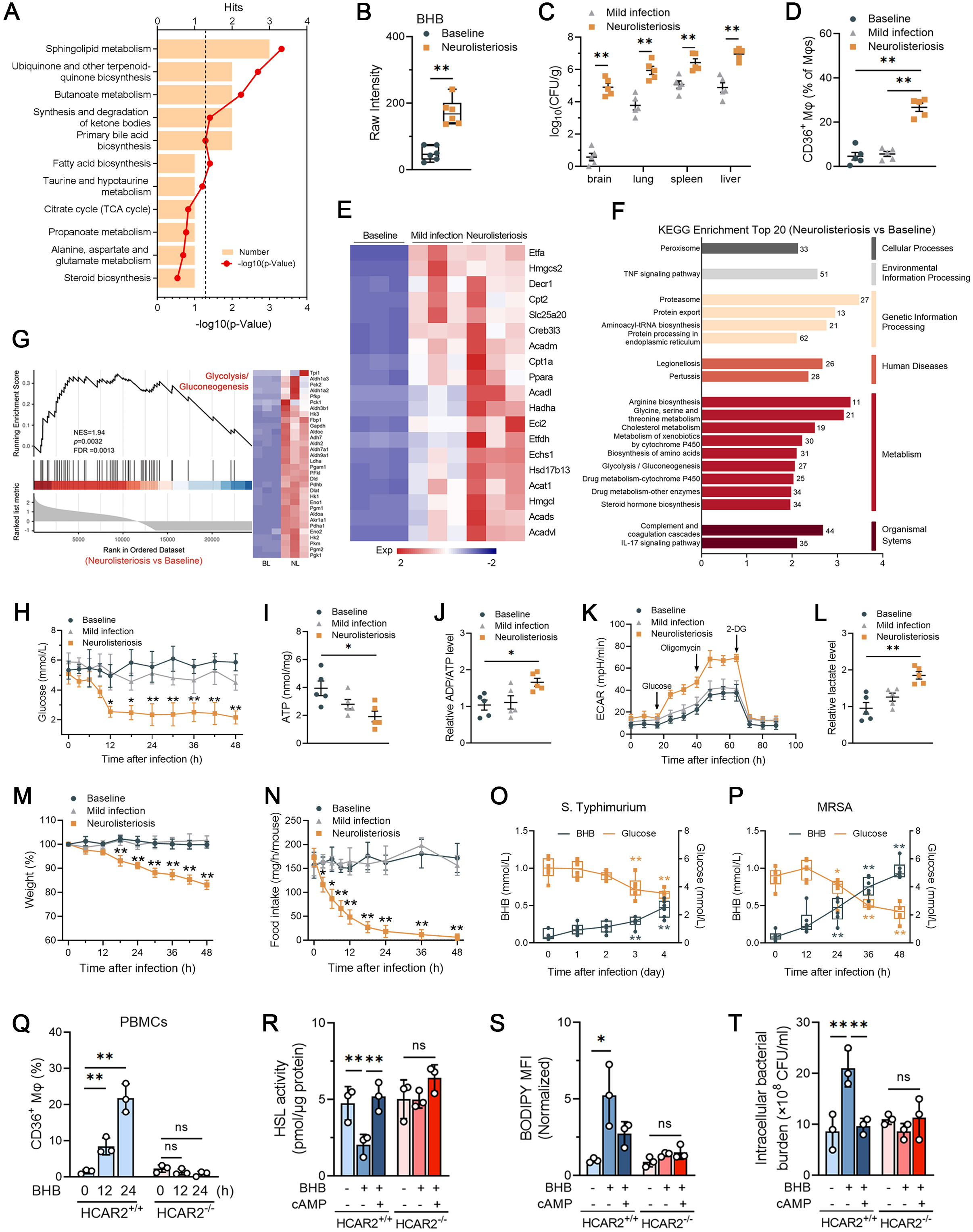
Additional results of β-hydroxybutyrate promote bacterial neuroinvasion, related to Figure 4. (A) KEGG enrichment analysis of lipid metabolites. The horizontal axis represents the number of -log_10_(p-Value), and the dashed line represents p-Value=0.05. Five of the top six enrichment pathways are above the significance level, namely Sphingolipid metabolism (Sphingosine, Sphinganine, Phytosphingosine. p-Value= 0.0005), Ubiquinone and other terpenoid-quinone biosynthesis (Vitamin K2, Vitamin K1. p-Value= 0.002), Butanoate metabolism [3-hydroxybutyrate (BHB), Succinic acid. p-Value=0.006], Synthesis and degradation of ketone bodies [3-hydroxybutyrate (BHB), Acetyl coenzyme A. p-Value=0.04], Fatty acid biosynthesis (Dodecanoic acid. p-Value=0.04). (B) Mass spectrometry analysis of serum BHB levels. (C-N) Mice in the mild infection group were intraperitoneally infected with 2×10^6^ CFU, whereas the neurolisteriosis group was intraperitoneally infected with 2×10^8^ CFU of *L. monocytogenes*. (n=5 for each group) (C-D) Bacterial burdens in the organs (C) and percentages of CD36^+^ macrophage (D) were detected at 24 hours post infection. (E-G) Transcriptomics analysis of liver tissues from mice (n=3 for each group). (E) Heatmap of differentially expressed (DE) genes related to ketogenesis. (F) Top 20 KEGG enrichment analysis of function of DE genes in neurolisteriosis group and baseline group. (G) Gene-set enrichment analysis for glycolysis/gluconeogenesis in neurolisteriosis group and baseline group. (H) Biochemical analysis of serum glucose levels in mice during *L. monocytogenes* infection. (I-L) Levels of ATP generation (I), ADP/ATP ratios (J), seahorse analysis of the ECAR (K) and relative abundances of lactate (L) in liver tissues of mice were detected at 24 hours (unless otherwise indicated) post infection. (M-N) Changes in body weight (M) and food intake (N) of the mice post infection. (O-P) Biochemical analysis of serum BHB and glucose levels in mice intraperitoneally infected with 1×10^8^ CFU of *S. typhimurium* (O) and 2×10^8^ CFU of MRSA (P). (n=5 for each group) (Q) PBMCs were isolated form HCAR2^+/+^ and HCAR2^-/-^ mice, and then treated with BHB (10 mM) for 24 hours. Percentages of CD36^+^ macrophages were analyzed by flow cytometry. (R-T) HCAR2^+/+^ and HCAR2^-/-^ macrophages were pretreated with BHB (10 mM) or cAMP (1 mM) for 24 hours and then infected with *L. monocytogenes* (MOI=10) for 6 hours. HSL activity (R), normalized MFI of BODIPY (S) and the penetrating bacterial burden in the lower chamber of Transwell system (T) were quantified. Experiments were performed in duplicate (C-P) or triplicate (Q-T). Data are presented as mean ± SEM [n=3 in (Q-T), n=5 in (B-D, H-P)]. *p < 0.05, **p < 0.01. Abbreviations: BL, baseline; NL, neurolisteriosis; ns, nonsignificant; MFI, mean fluorescence intensity; HSL, hormone-sensitive lipase.

**Figure S8.**
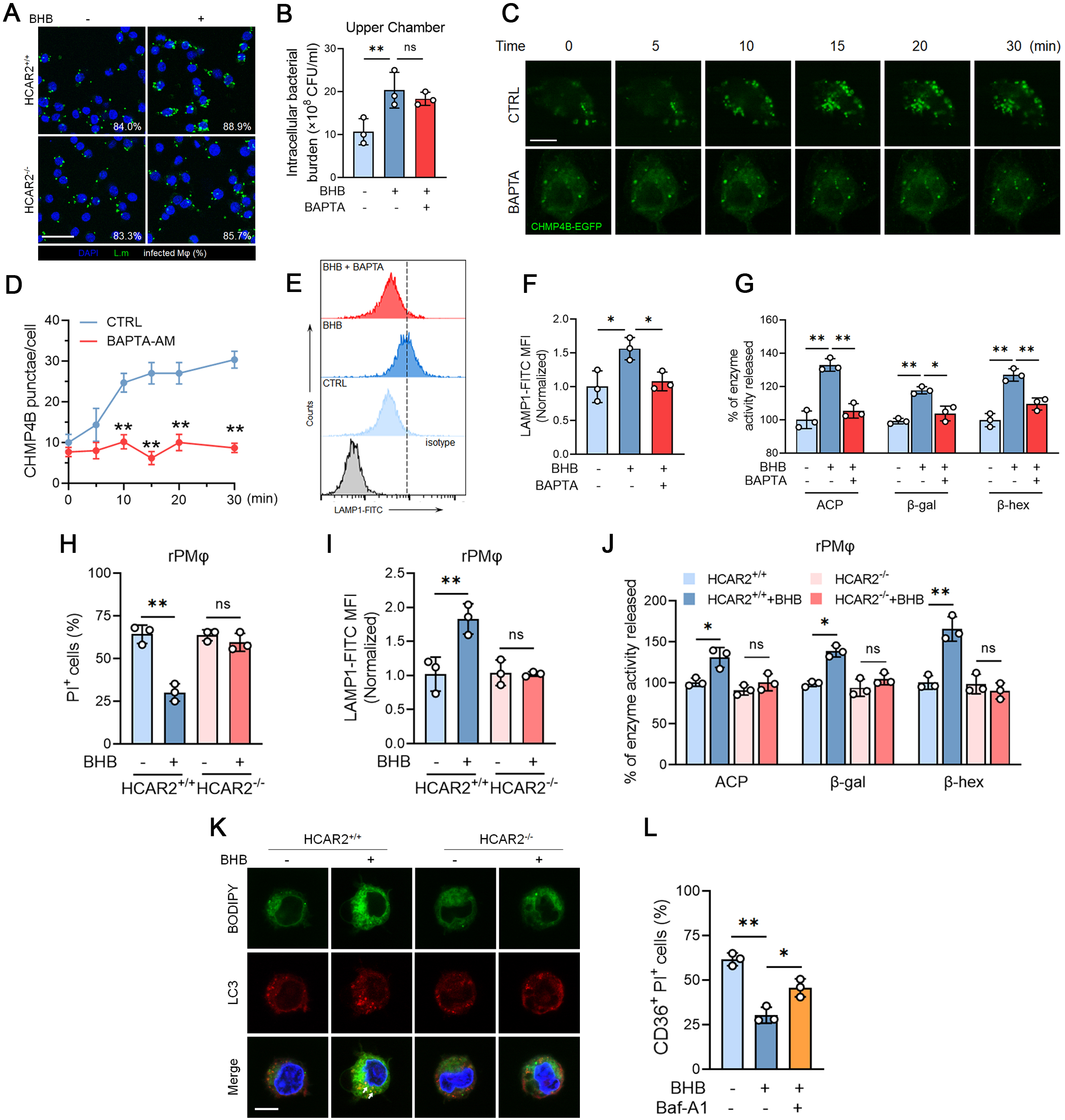
Additional results on the role of β-hydroxybutyrate in regulating macrophage survival, related to Figure 5. (A) Immunofluorescence staining of infected macrophages. The HCAR2^+/+^ and HCAR2^-/-^ macrophages were isolated and infected with *L. monocytogenes* (MOI=10) for 6 hours, and then cells were washed by PBS containing gentamicin. Intracellular bacteria (green) in macrophages were stained afterwards. Scale bar, 10 μm. (B-G) Primary peritoneal macrophages were extracted from HCAR2^+/+^ and HCAR2^-/-^mice and then treated with BHB (10 mM) or BAPTA (10 μM) 24 hours before infection (MOI=10, 6 hours). (B) Intracellular bacterial burdens of macrophages in the upper chamber of Transwell system. (C) Time-lapse images of CHMP4B-EGFP in macrophages were collected every 5 minutes post BHB treatment with confocal microscopy. (D) Quantification of CHMP4B-EGFP puncta. Flow cytometry analysis (E) and normalized MFI (F) of LAMP1 expression on the cell surface. (G) Quantification of the activities of lysosomal enzymes released into the culture medium. (H-J) Resident peritoneal macrophages were isolated and then infected with *L. monocytogenes* (MOI=10) for 6 hours. Percentage of PI^+^ cells (H) and the normalized MFI of LAMP1 expression on the cell surface (I) were assessed by flow cytometry. (J) Quantification of the activities of lysosomal enzymes released into the culture medium. (K) Primary peritoneal macrophages were isolated from HCAR2^+/+^ and HCAR2^-/-^mice and then infected with *L. monocytogenes* (MOI=10) for 6 hours. The colocalization of BODIPY (green) and LC3 (red) in infected macrophages was examined by immunofluorescence. Scale bar, 10 μm. (L) Percentage of CD36^+^PI^+^ cells at 24 hours post infection was quantified by flow cytometry. CD36^+^ macrophages were treated with BHB (10 mM) 24 hours before infection. Baf-A1 (100 nM) was added into the culture medium 6 hours before BHB stimulation. Experiments were performed in triplicate. Data are presented as mean ± SEM (n=3). *p < 0.05, **p < 0.01. Abbreviations: ns, nonsignificant; BHB, β-hydroxybutyrate; MFI, mean fluorescence intensity; rPMφs, resident peritoneal macrophages.

**Figure S9.**
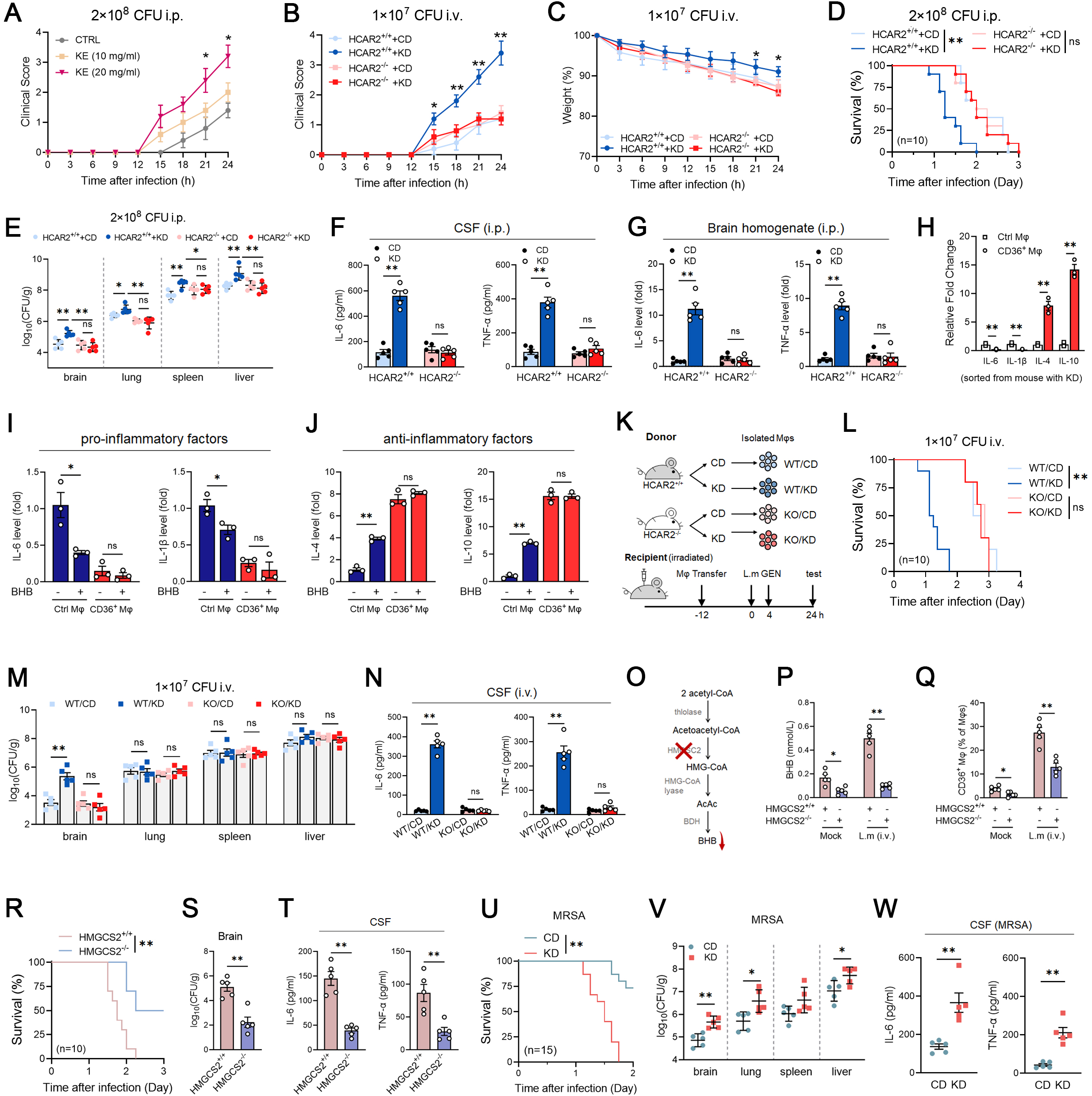
Additional results of ketogenesis increases bacterial neuroinvasion, related to Figure 6. (A) Clinical scores of mice supplemented with KE as shown in Figure 6A. (B-C) Clinical scores (B) and changes in body weight (C) of mice in the ketogenic diet experiments as shown in Figure 6F. (D-G) HCAR2^+/+^ and HCAR2^-/-^ mice were treated with ketogenic or control diets, respectively, followed by intraperitoneal infection of 2×10^8^ CFU *L. monocytogenes*. (n=10 for each group) (D) Survival of mice exposed to *L. monocytogenes*. (E) Bacterial burdens in the organs. Pro-inflammatory factor levels in CSF (F) and brain homogenates (G). (H-J) MRNA expressions of inflammatory factors in control macrophages and CD36^+^macrophages sorted from peripheral blood of mice with ketogenic diets (H) or with BHB (10 mM) pretreatment (I-J). (K-N) (K) Schematic diagram of adoptive macrophage transfer experiments. Peritoneal macrophages isolated from HCAR2^-/-^ and HCAR2^+/+^ mice with ketogenic or control diets were injected into irradiated recipient mice. After recovery for 12 hours, recipient mice were intravenously injected with 1×10^6^ CFU of *L. monocytogenes.* Gentamicin (30 mg/kg) was administered intraperitoneally 4 hours after infection. (n=10 for each group) (L) Survival of recipient mice. (M) Bacterial burdens in organs. (N) Pro-inflammatory factor levels of recipient mice at 24 hours post infection. (O-T) (O) Anabolic mechanism of BHB. (P-T) HMGCS2^+/+^ and HMGCS2^-/-^ mice were intravenously infected with 1×10^7^ CFU of *L. monocytogenes*. Samples were collected for examination 24 hours post infection. (n=10 for each group) (P) Serum BHB levels. (Q) Percentages of CD36^+^ macrophages. (R-T) Survival rates of mice (R), bacterial burdens in the brain (S) and pro-inflammatory factor levels in the CSF (T) of infected mice. (U-W) Mice were treated with ketogenic or control diets, respectively, followed by intraperitoneal infection of 2×10^8^ CFU MRSA. (n=15 for each group) (U) Survival of mice exposed to MRSA. (V) Bacterial burdens in the organs (brain, lung, spleen, and liver). (W) Pro-inflammatory factor levels in the CSF. Experiments were performed in duplicate (A-G, K-W) or triplicate (H-J). Data are presented as mean ± SEM [n=3 in (H-J), n=5 in (A-C, E-G, M-N, P-Q, R-T, U-V)]. *p < 0.05, **p < 0.01. Abbreviations: ns, nonsignificant; i.v., intravenous; i.p., intraperitoneal; PMφ, peritoneal macrophages; ctrl, control CD36^+^ Mφ, CD36^+^ macrophages; GEN, gentamicin; CD, control diet; KD, ketogenic diet.

**Figure S10.**
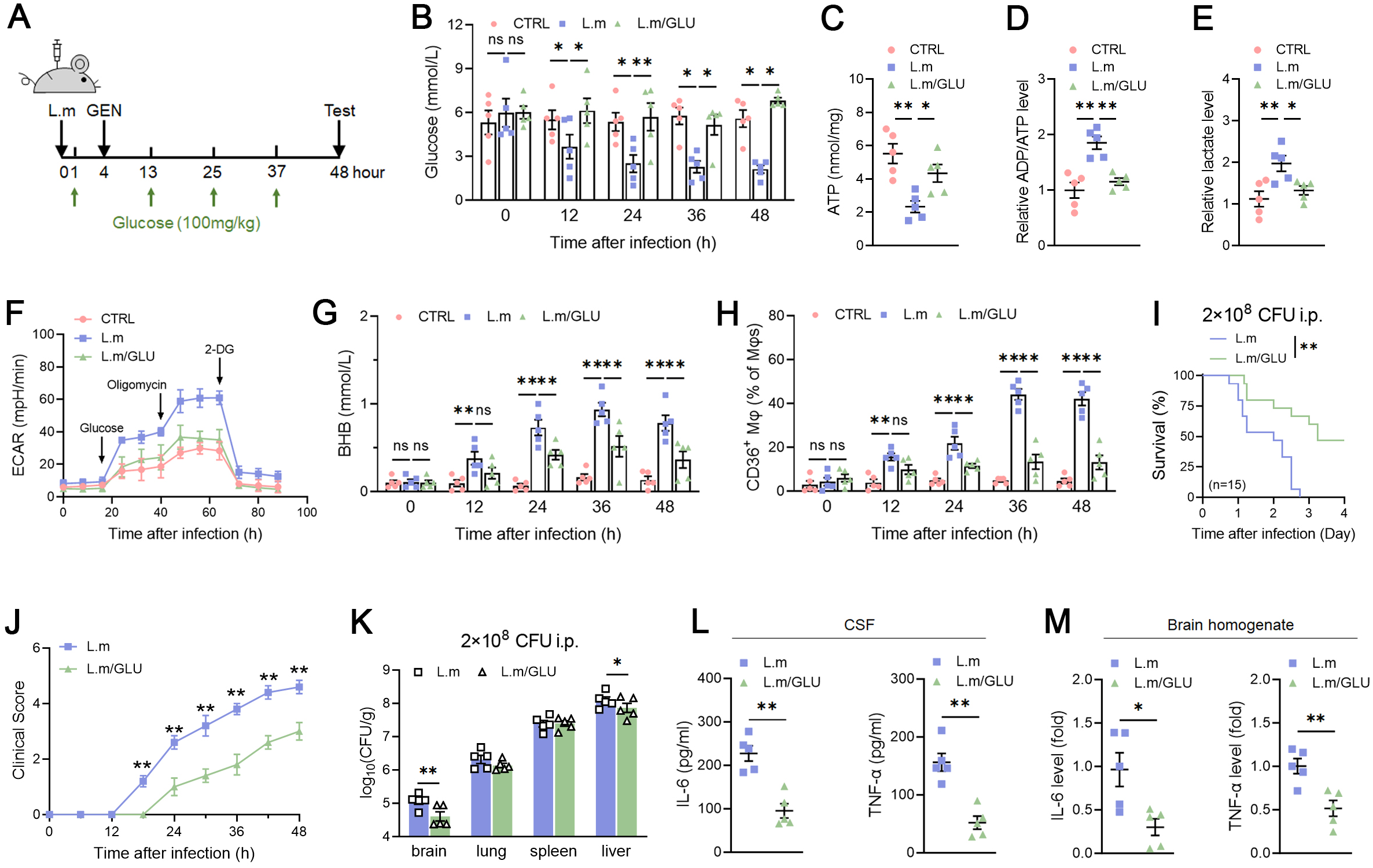
Increased glucose levels alleviate bacterial neuroinvasion. (A-M) (A) Schematic diagram of *in vivo* glucose supplementation. C57BL/6 mice were intraperitoneally challenged with 2×10^8^ CFU *L. monocytogenes*, and glucose (100 mg/kg) was injected 1 hour post infection, with repeated injections every 12 hours. Samples were collected at 48 hours (unless otherwise indicated) post infection. (n=15 for each group) (B) Serum glucose levels. (C-F) Levels of ATP generation (C), ADP/ATP ratios (D), relative abundances of lactate (E) and seahorse analysis of the ECAR (F) in liver tissues of mice at 48 hours post infection. (G) BHB levels. (H) Percentages of CD36^+^ macrophages in the PBMCs of infected mice. (I) Survival of mice exposed to *L. monocytogenes* with or without glucose supplementation. (J) Clinical scores. (K) Bacterial burdens in parenchymal organs. (L-M) Pro-inflammatory factor levels in CSF (L) and brain homogenates (M) of mice at 48 hours post infection. Experiments were performed in duplicate. Data are presented as mean ± SEM (n=5). *p < 0.05, **p < 0.01. Abbreviations: ns, nonsignificant; GEN, gentamicin; GLU, glucose.

**Figure S11.**
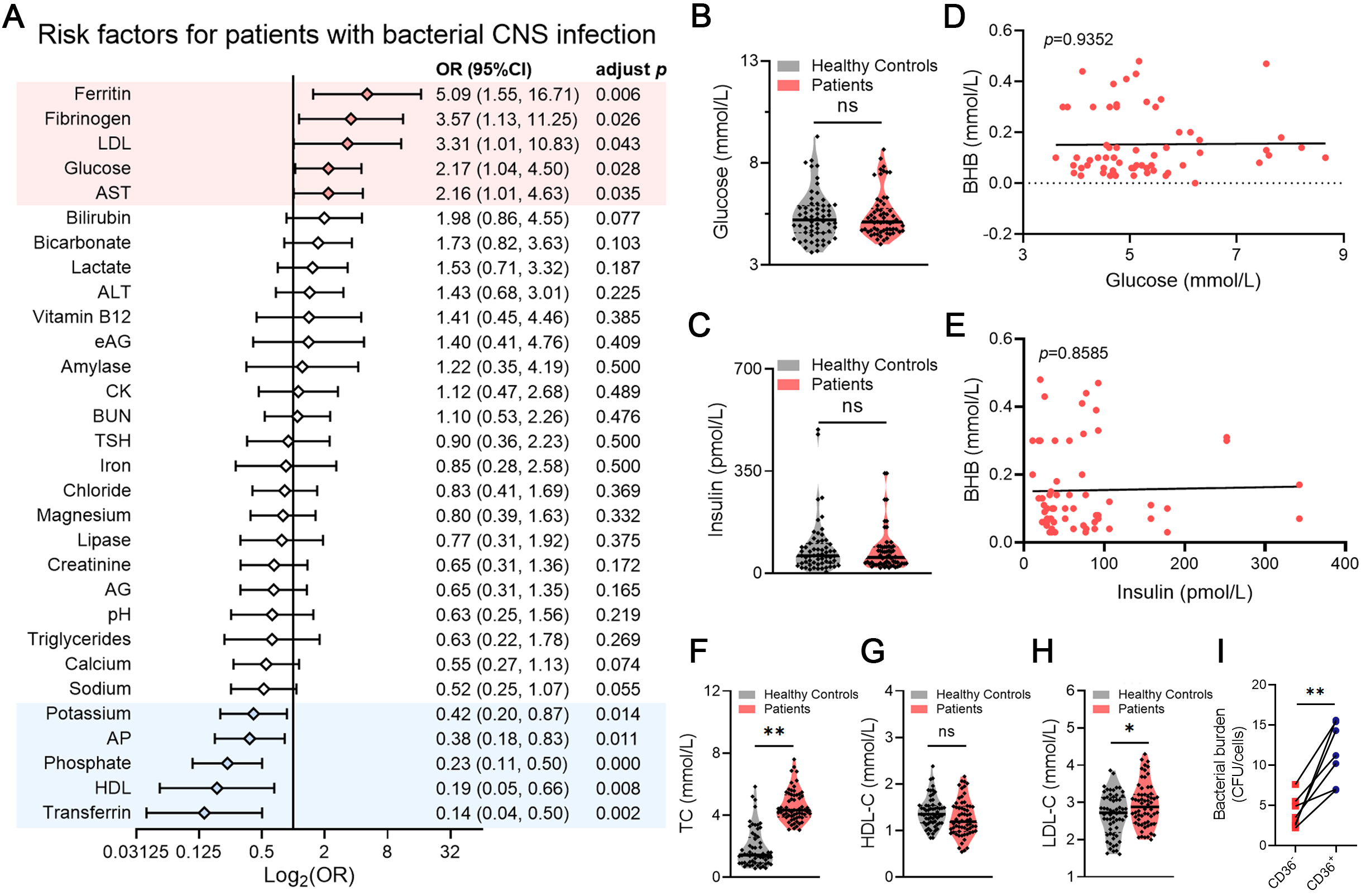
Additional results of high levels of β-hydroxybutyrate associate with bacterial neuroinvasion in humans, related to Figure 7. (A) Risk factors for patients with bacterial CNS infection in the MIMIC IV database. (B-E) Glucose (B) and insulin (C) levels in the serum from recruited healthy controls and bacterial neuroinvasion patients. Correlation between serum glucose levels and BHB levels (D), and correlation between serum insulin levels and BHB levels (E) in bacterial neuroinvasion patients were calculated. R, correlation coefficient. (F-H) Serum triglyceride (TC) (F), high density lipoprotein (HDL-C) (G), and low density lipoprotein (LDL-C) (H) from healthy controls and patients. (I) The intracellular bacterial burdens of CD36^+^ and CD36^-^ macrophages sorted from peripheral blood of patients by were examined by flow cytometry. Data are presented as mean ± SEM. *p < 0.05, **p < 0.01. Abbreviations: ns, nonsignificant.

